# Rapid and parallel adaptive mutations in spike S1 drive clade success in SARS-CoV-2

**DOI:** 10.1101/2021.09.11.459844

**Authors:** Kathryn E. Kistler, John Huddleston, Trevor Bedford

## Abstract

Given the importance of variant SARS-CoV-2 viruses with altered receptor-binding or antigenic phenotypes, we sought to quantify the degree to which adaptive evolution is driving accumulation of mutations in the SARS-CoV-2 genome. Here we assessed adaptive evolution across genes in the SARS-CoV-2 genome by correlating clade growth with mutation accumulation as well as by comparing rates of nonsynonymous to synonymous divergence, clustering of mutations across the SARS-CoV-2 phylogeny and degree of convergent evolution of individual mutations. We find that spike S1 is the focus of adaptive evolution, but also identify positively-selected mutations in other genes that are sculpting the evolutionary trajectory of SARS-CoV-2. Adaptive changes in S1 accumulated rapidly, resulting in a remarkably high ratio of nonsynonymous to synonymous divergence that is 2.5X greater than that observed in HA1 at the beginning of the 2009 H1N1 pandemic.

## Introduction

After 20 months of global circulation, basal lineages of SARS-CoV-2 have been almost completely replaced by derived, variant lineages. These lineages are classified by the WHO as variants of concern (VOCs) or variants of interest (VOIs) based on genetic, phenotypic and epidemiological differences [1]. The effort to track the spread of these variants (and of the pandemic in general) through genomic epidemiology has resulted in a massive corpus of sequenced viral genomes. In the GISAID EpiCoV database alone, there are 2.5 million sequences and counting as of the end of July 2021 [2]. This thorough sampling offers an opportunity to investigate the evolutionary dynamics of a virus as it entered a naive population, spread rampantly, and, subsequently, began to transmit through previously exposed hosts. Here, we are particularly interested in whether SARS-CoV-2 viruses show phylogenetic evidence of adaptive evolution during the first year and a half of transmission in humans.

Seasonal influenza and seasonal coronaviruses both exhibit continual adaptive evolution during endemic circulation in the human population. In the case of influenza H3N2, transmission through an exposed host population results in adaptive evolution within hemagglutinin (HA). The HA1 subunit of hemagglutinin both mediates binding to host cell receptors and is the primary target for neutralizing antibodies. Thus, in the context of an exposed host, selection for receptor binding avidity [3] and for escape from humoral immunity [4] drive fixation of mutations in the HA1 subunit. The coronavirus protein subunit equivalent in function to HA1 is spike S1. Previously, we showed that at least two seasonal coronaviruses (229E and OC43) exhibit adaptive evolution concentrated in the S1 subunit of spike [5]. By demonstrating that strong immune responses to a particular historical isolate of 229E do not neutralize 229E viruses that circulate years afterwards, Eguia et al confirmed that 229E evolves antigenically [6].

Standard methods used to detect adaptive evolution in seasonal influenza and seasonal coronaviruses rely on the fixation (or near fixation) of nonsynonymous changes, and thus require years or decades of evolutionary time. These methods are ill-fit to identify early adaptive evolution of a virus that has experienced a recent spillover event. For example, the common ancestor of globally circulating SARS-CoV-2 viruses is currently no earlier than January 2020, corresponding to the base of clade 20A or lineage B.1 (nextstrain.org/ncov/gisaid/global). Here, we present a new method to identify regions of the genome undergoing adaptive evolution, which is well-suited to early time points. This method correlates clade success with the accumulation of protein-coding changes in certain genes. We apply this method to SARS-CoV-2 genomic data from Dec 2019 to May 2021, focusing on the period of VOC and VOI emergence.

With this method, we aim to present a rigorous quantification of the evolutionary process during this time and to show that the observed success of variant viruses is a result of adaptive, not neutral, evolution. We conduct these analyses across the SARS-CoV-2 genome to identify foci of adaptive evolution. We complement these results with analyses of *d*_*N*_*/d*_*S*_ accumulation, evolutionary dynamics, and convergent evolution to provide evidence that genetic changes are contributing to viral fitness and identify genomic regions that are responsible.

## Results

### Accumulation of nonsynonymous mutations in spike S1 correlates with clade success

RNA viruses are known for their remarkably high error rates and, thus, the rapid generation of mutations. Despite possessing some proof-reading capacity (a relatively rare function for an RNA virus), SARS-CoV-2 has been accumulating roughly 24-25 substitutions per year (nextstrain.org/groups/blab/ncov/adaptive-evolution/2021-05-15?l=clock, [7]). The null hypothesis is that these substitutions reflect neutral evolution: the result of genetic drift acting on random mutations. To determine whether this is true, or whether adaptive evolution is also contributing to the accumulation of mutations, we started by comparing substitution rates in different regions of the genome.

We built a time-resolved phylogeny with a balanced geographic and temporal distribution of samples collected between December 2019 and May 15, 2021 that includes 9544 viruses (Figure S1). For every internal branch on the phylogeny, we tallied the total number of mutations that occurred between the phylogeny root and that branch. We grouped deletion events with nonsynonymous single nucleotide polymorphisms (SNPs), as they are protein-changing and contribute to the evolution of some regions of the genome (Figure S2). Plotting mutation counts over time shows that spike S1 accumulates nonsynonymous changes at a rate of 8.4 × 10^*−*3^ substitutions/codon/year, or about 5.5 substitutions per year (Figure 1A). This is a disproportionate percentage of the genome-wide estimate of 24 substitutions per year. As a control, we counted S1 synonymous mutations, and found they accumulate at 2.0 × 10^*−*4^ substitutions/codon/year, close to the naive expectation from base composition that 22% of mutations should be synonymous. The per-codon rate of nonsynonymous mutation in S1 is roughly 17 times higher than in the RNA-dependent RNA polymerase (RdRp) gene.

**Figure 1.**
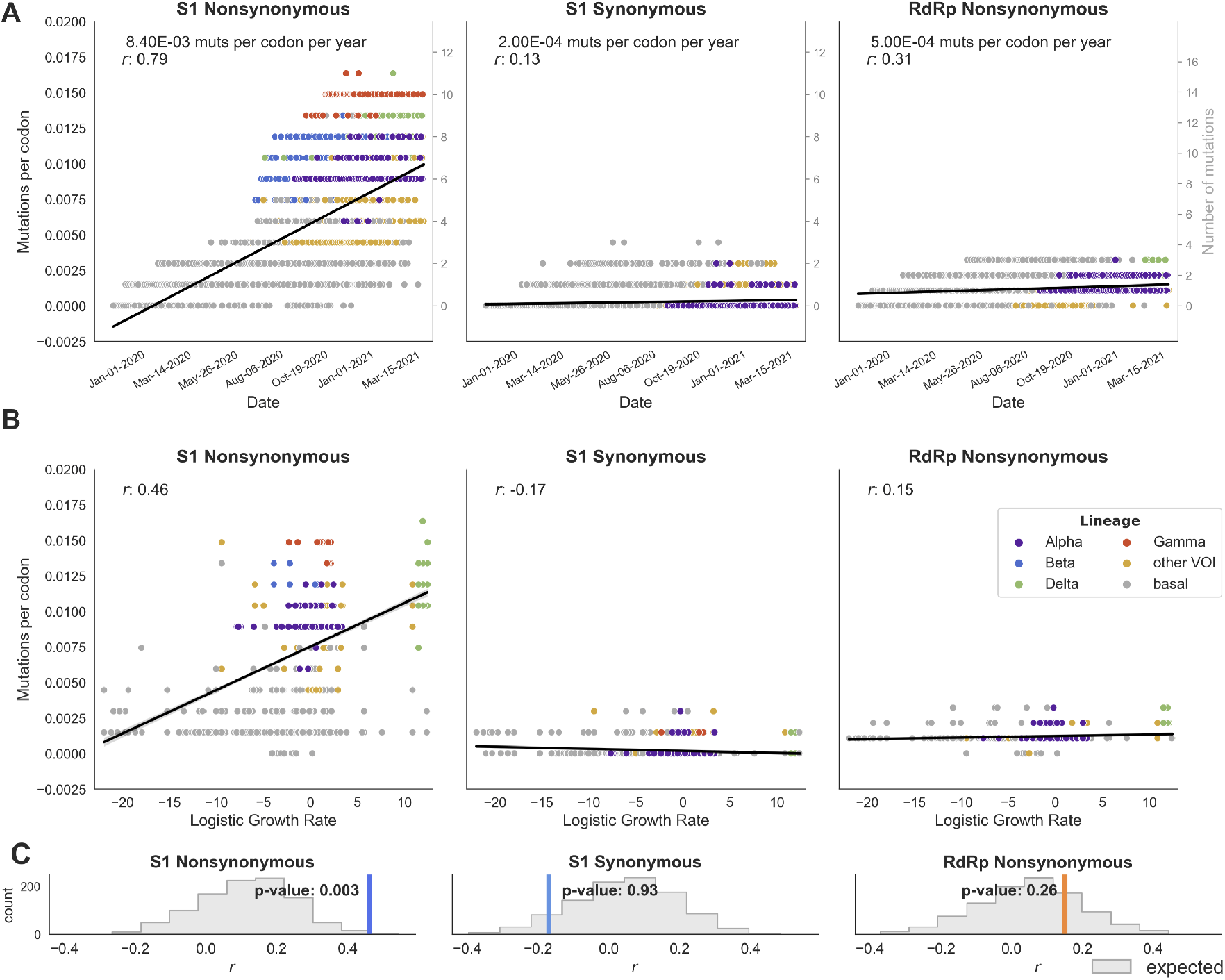
Accumulation of nonsynonymous S1 mutations is correlated with clade success. **A)** For every clade in the phylogeny, mutations relative to the root of the phylogeny are tallied and plotted against the date of the base of that clade. Nonsynonymous S1, synonymous S1, and nonsynonymous RdRp mutations are plotted separately. Nonsynonymous mutations include nonsynonymous SNPs and deletions. The primary axis (left, black ticks) displays mutations per codon, and the secondary axis (right, gray ticks) shows the absolute number of mutations accumulated in each clade. Each point is colored according to the lineage it belongs to. Points are fit by linear regression. **B)** For every clade, mutation accumulation (as in A) is plotted against logistic growth rate and the points are fit by linear regression. **C)** The empirical correlation coefficient *r* between mutation accumulation and logistic growth rate (colored bar) is compared to an expected distribution (gray) to yield a *p*-value. Expected values of *r* are determined from randomizing mutations across the phylogeny using a multinomial draw with mutation likelihood proportional to relative branch length. The results of 1000 iterations are shown.

We hypothesize that adaptive evolution is driving the high rate of S1 nonsynonymous substitutions relative to S1 synonymous substitutions and RdRp nonsynonymous substitutions. And, though each S1 substitution will have a different effect on fitness, this observation suggests that this class of mutations is, on average, under positive selection. If this is the case, we would expect a correlation between S1 substitutions and a clade’s evolutionary success: clades that happened to accumulate more S1 substitutions should have, on average, higher fitness (and hence faster growth in frequency) than clades that have accumulated fewer S1 substitutions. Based on this logic, we introduce a new method for detecting adaptive evolution, which looks for regions of the genome where mutation accumulation is associated with clade frequency growth. Because positive selection causes alleles or clades to increase in frequency in a logistic (rather than linear) fashion, we measure logistic growth rate and plot this versus mutation accumulation.

Clade success and the number of nonsynonymous S1 mutations are positively correlated, with a correlation coefficient *r* of 0.46 (Figure 1B). To test whether this correlation is greater than expected, we randomized the placement of mutations across branches of the phylogeny and computed a *p*-value between the empirical *r* and the distribution of *r* values from 1000 randomizations. The positive correlation between S1 mutations and logistic growth rate is statistically significant compared to the expected distribution (*p*=0.003), but is absent for S1 synonymous mutations and is not significant for RdRp substitutions (*p*=0.256) (Figure 1C).

We applied this method to every gene in the SARS-CoV-2 genome (Table 1 and Figure S3). The highest nonsynonymous mutation rate is observed in ORF8. However, ORF8 substitutions are not correlated with clade success (Table 1), and many lineages acquire premature stop codons in ORF8, indicating that the high rate of ORF8 substitutions is likely due, at least in part, to a lack of functional constraints. Mutations within other regions of the genome, including spike S2 and nucleocapsid (N), also accumulate at reasonably-high levels but do not correlate well with clade success (Table Table 1). Besides S1, only Nsp6 (*r*=0.35, *p*=0.011) and ORF7a (*r*=0.43, *p<*0.001) have a strong correlation with clade growth rates (Table 1, Figures S3 and S4).

**Table 1.** Genome-wide correlation between nonsynonymous mutation accumulation and logistic growth rate showing evolutionary rate of nonsynonymous substitutions (and deletions) per codon per year, the correlation coefficient (*r*) of mutation accumulation with logistic growth rate and the p-value of this correlation.

| Gene | Nonsyn Evolution rate<br>(subs/codon/year) | $r$ | p-value |
| --- | --- | --- | --- |
| Nsp1 | $0.1 \times 10^{-3}$ | 0.05 | 0.431 |
| Nsp2 | $0.2 \times 10^{-3}$ | -0.10 | 0.881 |
| Nsp3 | $1.3 \times 10^{-3}$ | 0.30 | 0.083 |
| Nsp4 | $0.8 \times 10^{-3}$ | 0.28 | 0.057 |
| Nsp5 | $0.5 \times 10^{-3}$ | -0.09 | 0.833 |
| Nsp6 | $3.4 \times 10^{-3}$ | 0.35 | **0.011** |
| Nsp7 | $0.3 \times 10^{-3}$ | -0.42 | 0.998 |
| Nsp8 | $0.3 \times 10^{-3}$ | 0.04 | 0.443 |
| Nsp9 | $0.7 \times 10^{-3}$ | 0.06 | 0.299 |
| Nsp10 | $0.1 \times 10^{-3}$ | -0.05 | 0.807 |
| RdRp | $0.5 \times 10^{-3}$ | 0.15 | 0.256 |
| Nsp13 | $0.8 \times 10^{-3}$ | 0.20 | 0.153 |
| Nsp14 | $0.3 \times 10^{-3}$ | -0.03 | 0.698 |
| Nsp15 | $0.3 \times 10^{-3}$ | 0.15 | 0.230 |
| Nsp16 | $0.3 \times 10^{-3}$ | -0.22 | 0.963 |
| S1 | $8.4 \times 10^{-3}$ | 0.46 | **0.003** |
| S2 | $3.5 \times 10^{-3}$ | 0.24 | 0.105 |
| ORF3a | $1.8 \times 10^{-3}$ | -0.06 | 0.865 |
| E | $1.6 \times 10^{-3}$ | 0.03 | 0.388 |
| M | $0.7 \times 10^{-3}$ | 0.29 | 0.045 |
| ORF6 | $0.2 \times 10^{-3}$ | 0.01 | 0.528 |
| ORF7a | $1.5 \times 10^{-3}$ | 0.43 | **<0.001** |
| ORF7b | $1.7 \times 10^{-3}$ | 0.22 | 0.050 |
| ORF8 | $16.5 \times 10^{-3}$ | 0.19 | 0.185 |
| N | $6.10 \times 10^{-3}$ | 0.21 | 0.222 |
| ORF9b | $2.10 \times 10^{-3}$ | 0.27 | 0.050 |

Though ORF7a substitutions appear highly correlated with clade success, this correlation is driven solely by the rapidly growing Delta variant, which possesses 3 mutations in ORF7a. Removing Delta clades from the analysis drops the *r* for ORF7a from 0.43 to 0.16, whereas *r* for S1 and Nsp6 only dip from 0.46 to 0.41, and from 0.35 to 0.32, respectively. This indicates that the correlation between S1 and Nsp6 substitutions and clade success is a general feature of SARS-CoV-2 lineages. Thus, the metric presented here provides evidence that SARS-CoV-2 is evolving adaptively and that the predominant locus of this evolution is spike S1.

### The ratio of nonsynonymous to synonymous divergence is highest in S1

A classical method for assessing the average directionality of natural selection on some region of the genome is *d*_*N*_*/d*_*S*_, measuring the divergence of nonsynonymous sites relative to synonymous sites. A *d*_*N*_*/d*_*S*_ value less than 1 indicates that the region is, on average, under purifying selection, while *d*_*N*_*/d*_*S*_ greater than 1 indicates positive selection on the region. Because even the most rapidly evolving genes are still subject to structural and functional constraints, it is rare for an entire gene to have a *d*_*N*_*/d*_*S*_ ratio greater than 1. For instance, the HA1 subunit of H3N2, which is the prototypical example of an adaptively-evolving viral protein, has *d*_*N*_*/d*_*S*_ of 0.37 [8].

For various regions of the SARS-CoV-2 genome, we computed the accumulation of nonsynonymous and synonymous divergence in 2-month windows between January 1, 2020 and May 15, 2021 (Figure 2A). This measures *d*_*N*_*/d*_*S*_ of branches leading to tips sampled within each 2-month window and captures the progressive enrichment of mutations by natural selection, i.e. mutations that persist and contribute the viral population will be captured in this measure, while mutations that die out will be excluded. The *d*_*N*_*/d*_*S*_ ratio within RdRp, S2, and the structural proteins Envelope (E), Membrane (M), and Nucleocapsid (N) is consistently under 1 at all timepoints (Figure 2B). However, *d*_*N*_*/d*_*S*_ within S1 increases over time, with an apparent inflection point in mid-2020, and the *d*_*N*_*/d*_*S*_ ratio exceeding 1 in late-2020 and 2021 with the most recent time point measured at 1.80. For comparison, we used the same methodology to compute *d*_*N*_*/d*_*S*_ for influenza H3N2, influenza H1N1pdm and seasonal coronavirus OC43 from 2009 to 2021 (Figure S5). We observe that following emergence in humans in 2009 of influenza H1N1pdm, *d*_*N*_*/d*_*S*_ in HA1 subunit peaked at 0.72 roughly a year after the beginning of that pandemic and declined in the following 4 years, while endemic viruses H3N2 and OC43 showed relatively stable *d*_*N*_*/d*_*S*_ over this same time period.

**Figure 2.**
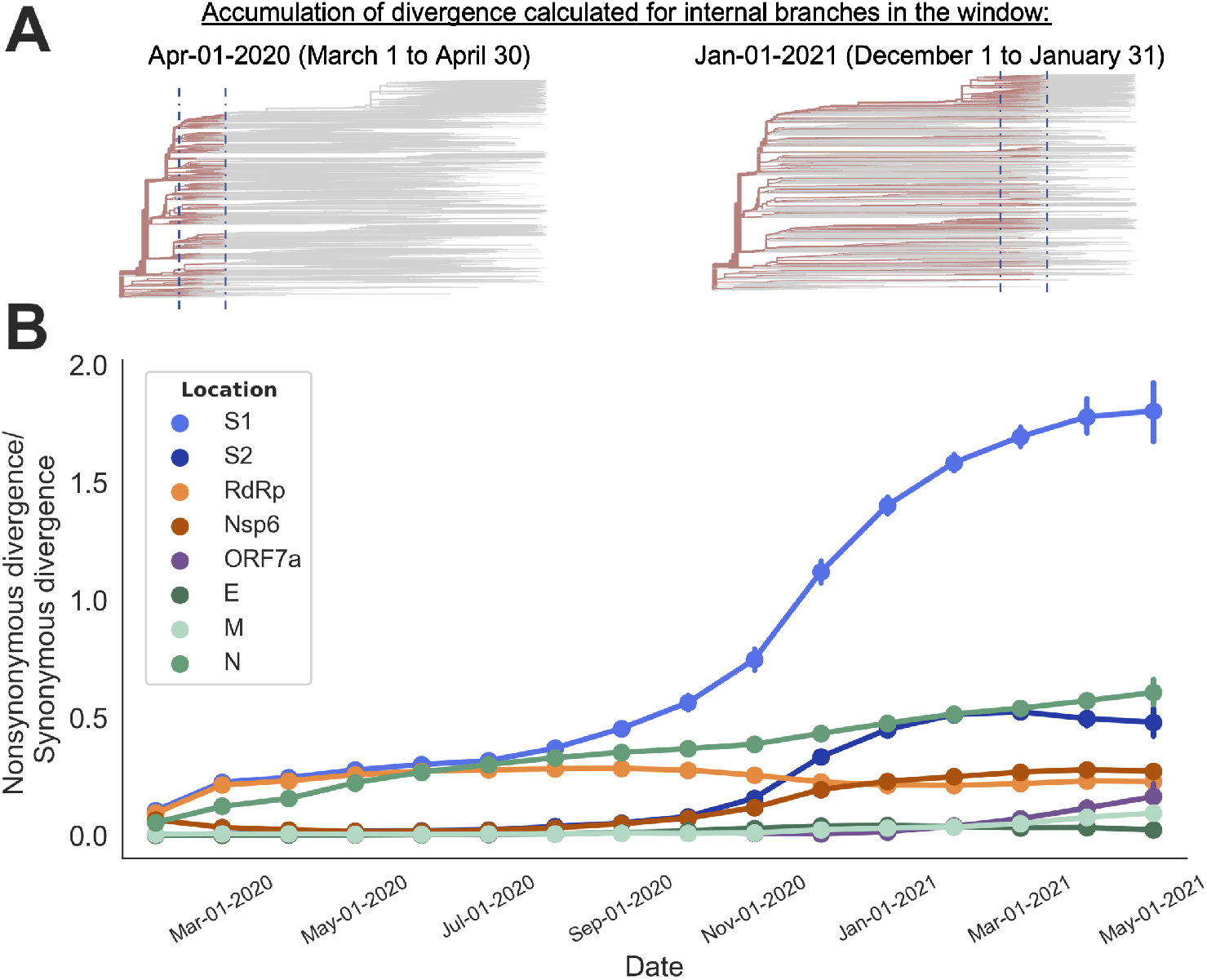
Ratio of nonsynonymous to synonymous divergence is highest S1. The ratio of nonsynonymous to synonymous divergence accumulation within various coding regions is calculated over time. The phylogeny of 9544 viral sequences is divided into overlapping 2-month windows between January 1, 2020 and May 15, 2021. **A)** The phylogeny is colored to indicate the paths on which divergence accumulation is calculated for two of these time windows. The dashed blue lines indicate the time windows centered at April 1, 2020 (left) and January 1, 2021 (right). Every internal branch within these windows and the phylogenetic path that connects that branch back to the root is highlighted in red. The accumulation of divergence is calculated along these paths. Nonsynonymous divergence is calculated as the nonsynonymous Hamming distance between the sequence of an internal branch and the root sequence, normalized by the total possible number of nonsynonymous sites. The same is done for synonymous divergence. **B)** The ratio of nonsynonymous to synonymous is calculated for various coding regions within the genome. Each point shows the mean and 95% confidence interval of this ratio for all internal branches present in a 2-month window (centered at the date indicated on the x-axis).

The increase over time in SARS-CoV-2 S1 *d*_*N*_*/d*_*S*_ could be due to a variety of reasons. Two non-mutually exclusive hypotheses include the appearance of a new selective pressure on S1 substitutions, or the acquisition of mutations that change the mutational landscape to be more permissive towards S1 substitutions. Regardless of the cause, this change suggests a temporal structure to the adaptive evolution in the S1 subunit of SARS-CoV-2.

### Nonsynonymous mutations in spike S1 cluster temporally

A hint of this temporal structure can be seen by tracing individual mutational paths through the tree, from root to tip. Figure S6 plots the accumulation of nonsynonymous S1 mutations along ten representative paths, leading to 10 different emerging lineages. Along each of these paths, there appears to be an initial period of relative quiescence, followed by a burst of S1 substitutions. To test whether this temporal clustering of mutations differs from what would be expected given the phylogenetic topology and the total number of observed S1 substitutions, we calculated wait times between mutations (diagrammed in Figure 3A). Briefly, we created a null expectation by running 1000 iterations of mutation randomization in which the phylogenetic placement of every observed mutation is shuffled. The distribution of wait times is dependent on tree topology and total number of mutations, so the expectation is different for each category of mutations (Figure S7).

**Figure 3.**
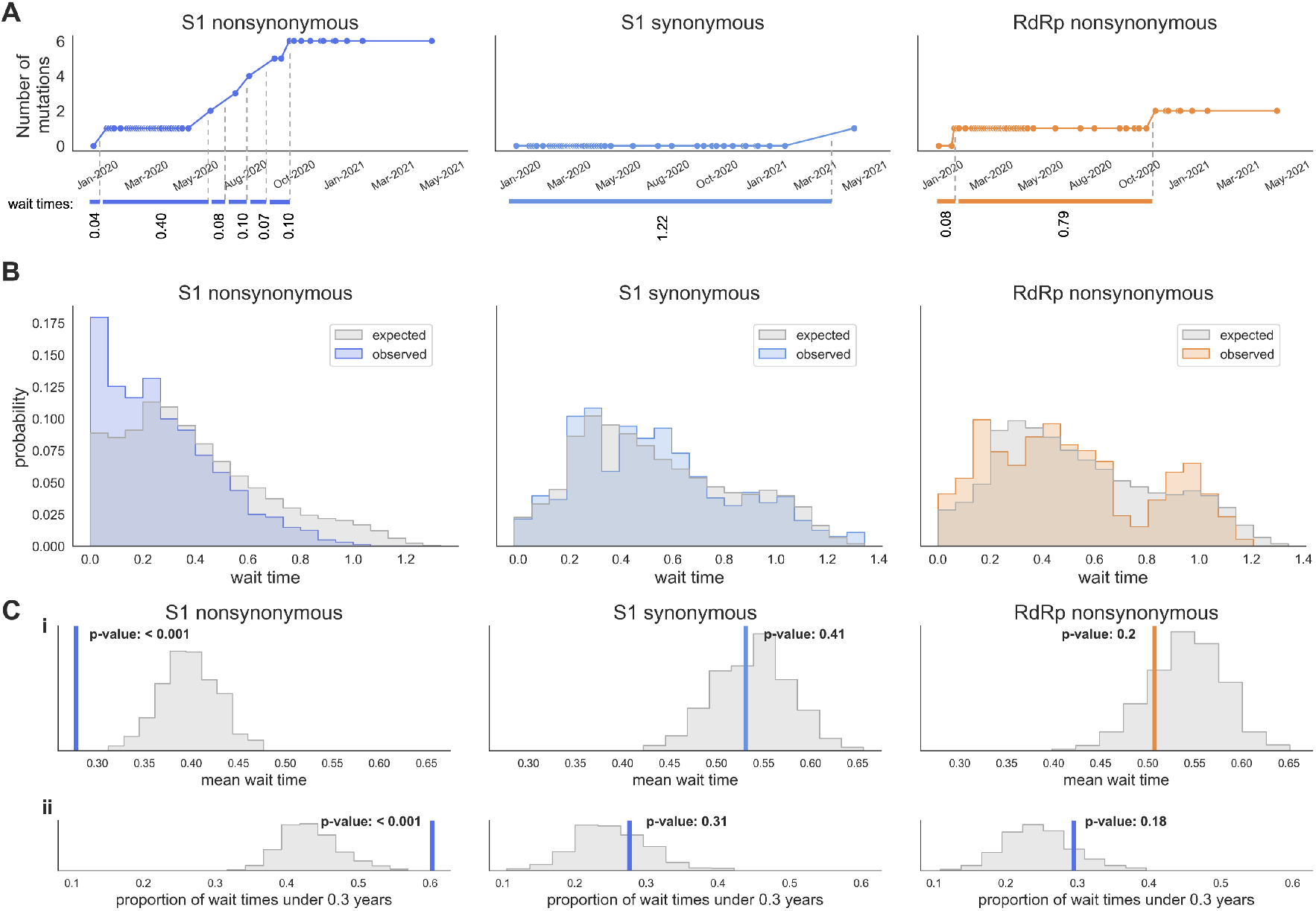
S1 substitutions are temporally clustered. **A)** Time line showing accumulation of S1 nonsynonymous, synonymous, and RdRp nonsynonymous mutations between the root and a representative tip (isolate USA/ME-HETL-J3202/2021) with wait times between mutations illustrated in below. The exact date of a mutation is chosen by randomly selecting a date along the branch the mutation occurs on. **B)** Distribution of wait times between S1 nonsynonymous, S1 synonymous, and RdRp nonsynonymous mutations. Empirical wait times (in color) are plotted along with expected wait times (gray). Expected wait times are determined from randomizing mutations across the phylogeny using a multinomial draw with mutation likelihood proportional to relative branch length. The results of 1000 iterations are shown. **C) i)** The mean empirical wait time from 1000 iterations of the analysis (colored bar) is compared to the distribution of mean expected wait times (gray) to yield a *p*-value. **ii)** The proportion of observed wait times under 0.3 years (colored bar) is compared to the distribution of expected wait times under 0.3 years (gray).

If mutations are clustered, there should be an excess of short wait times in the empirical data relative to the expectation. This is what we observe for S1 nonsynonymous mutations, where the distribution of wait times is left-skewed, with an overabundance of short wait times compared to the expected distribution (Figure 3B). The mean wait time between observed S1 substitutions is significantly lower than the expected mean wait time (*p<*0.001), while there is no significant difference for S1 synonymous or RdRp wait times (Figure 3Ci). This difference is driven by short wait times because there is a significant difference between the proportion of observed versus expected wait times under 0.3 years for S1 nonsynonymous, but not S1 synonymous or RdRp, mutations (Figure 3Cii). These results indicate a temporal structure to the adaptive evolution of SARS-CoV-2 within the S1 subunit, which is characterized by mutation clustering.

### Specific mutations associated with successful clades

We next sought to identify specific adaptive mutations throughout the genome. We note that convergent evolution is a good indicator of positive selection because each additional independent occurrence on the phylogeny of the mutation is increasingly unlikely under neutral evolution. As other groups have reported, there are many mutations shared by the VOCs that have arisen via convergent evolution [9–11]. Here, we combine this observation of convergent evolution with logistic growth rate to find mutations that have arisen in the SARS-CoV-2 population multiple, independent times and expand into successful clades after each occurrence.

In this analysis, we focus on the evolutionary dynamics of SARS-CoV-2 during the period of time between the emergence of this virus in humans and mid-May 2021. We estimate that, during this period of time, VOC viruses are primarily competing with basal SARS-CoV-2 viruses. This allows us to examine the overall fitness effects of specific mutations in viral lineages that are successful during this period of time. After May 2021, VOCs comprise a majority of the global virus population, and similar analyses on later time points would speak to the relative competitiveness of the variants.

For every deletion and substitution observed on the phylogeny, we tallied the number of independent occurrences and found the mean logistic growth rate of all clades where this mutation occurred. We limited this analysis to internal branches with 15 or more descending samples to limit the influence of stochasticity and sequencing errors that often occur on terminal branches. As expected, the bulk (84%) of mutations occur just once. Roughly 4% of mutations arose 4 or more times, and the majority of these mutations are located in S1 (Figure 4A). For seven of these convergently-evolved mutations, the mean growth rate is higher than the tree-wide average growth rate. For three of these mutations (S:95I, S:452R and ORF1a:3675-3677del), the mean growth rate exceeds the 90th percentile of mean growth rates expected from a mutation that occurs the same number of times on a randomized tree (Figure 4B).

**Figure 4.**
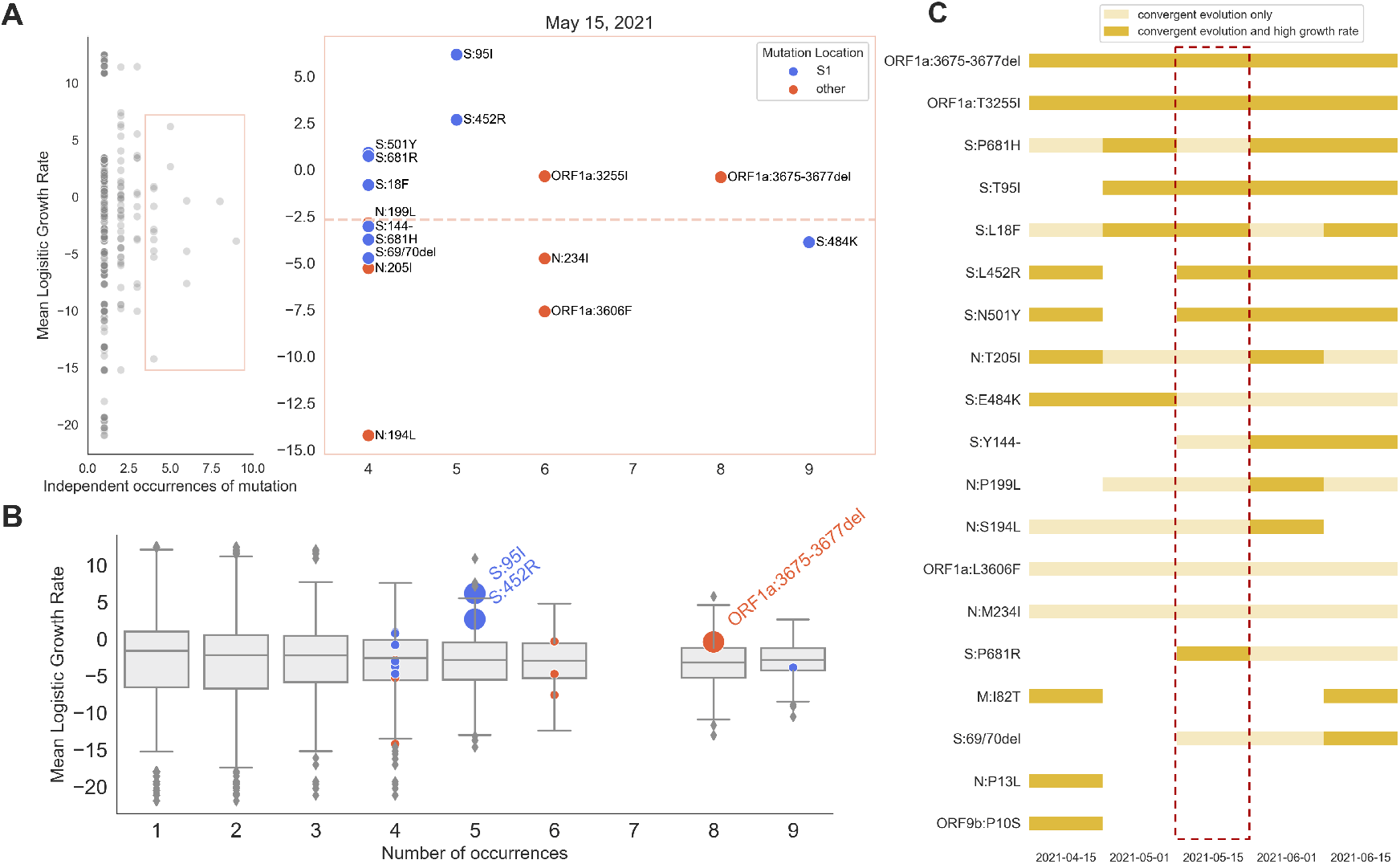
A 3-amino acid deletion in Nsp6 displays convergent evolution and occurs in successful clades. **A)** Every mutation observed on internal branches of the phylogeny is plotted according to the number of times this mutation occurs on the tree and the mean logistic growth rate of all clades it occurs in. Convergently-evolved mutations that appear 4 or more times across the phylogeny are shown in the inset. The average growth rate of all clades is shown with a dotted line. **B)** Observed mean growth rates of convergently-evolved mutations are compared to the mean growth rate expected for a mutation occurring on the phylogeny the same number of times. Convergently-evolved mutations that have a mean growth rate falling at or above the 90th percentile of the expected values are labeled. **C)** The analysis shown in panel A was completed for 5 time points, spanning two months. Each date represents the maximum date of sequences included in the analysis. Mutations that occur at least 4 times (convergent mutations) and result in a higher-than-average mean growth rate are shown in dark yellow. Mutations that display convergent evolution but do not result in high growth rates are in light yellow. The primary analysis was done at timepoint 2021-05-15 (outlined in red).

This analysis reveals influential mutations during a snapshot of time in the ongoing adaptive evolution of SARS-CoV-2. In mid-May 2021, the Delta variant was rising in frequency. Both S1 mutations we identified as important drivers of adaptive evolution (S:95I and S:452R) are present in the Delta variant as well as a handful of other emerging lineages (Figure S8). The specific mutations identified by this analysis will vary over time and depend on a multitude of factors (genetic, epidemiological, and otherwise) that determine clade success. However, ORF1a:3675-3677del consistently appears as a top hit (Figure 4C, and Figure S9). Remarkably, this deletion, which ablates amino acids 106-108 of Nsp6, arose 8 independent times and emerging lineages descend from each branch this deletion occurs on (Figure S8).

Because recombination is common in coronaviruses [12, 13], we investigated the possibility that these 8 occurrences of the ORF1a:3675-3677 deletion were due to recombination, rather than convergent evolution. We considered all pairs of lineages containing this mutation as potential recombinants and compared informative mutations in the potential donor and acceptor. The closest informative mutations flanking ORF1a:3675-3677del are not shared by any pairs of lineages, offering a lack of evidence for recombination and strong support for convergent evolution.

### A 3-amino acid deletion in Nsp6 is associated with accumulation of S1 substitutions

The ORF1a:3675-3677 deletion in Nsp6 exhibits striking convergent evolution and consistently precedes successful viral lineages. Because we have shown that S1 mutation accumulation is also associated with clade success, we next asked whether there is a relationship between the number of S1 substitutions in clades containing ORF1a:3675-3677del.

We created an expectation for the mean number of S1 mutations that should be observed in clades with ORF1a:3675-3677del by generating 100 randomized trees where the mutation occurred on 8 branches selected by a multinomial draw. To make the expectation as fair as possible, we constrained the randomized branches to be on or after the date that the first Nsp6 deletion was observed. Under this expectation, there is no difference between the mean number of S1 or RdRp substitutions in clades that have the ORF1a:3675-3677 deletion versus clades that do not (Figure 5A, left). However, in the empirical phylogeny, there are significantly more S1 substitutions in clades with the Nsp6 deletion versus clades without (Figure 5A, right).

**Figure 5.**
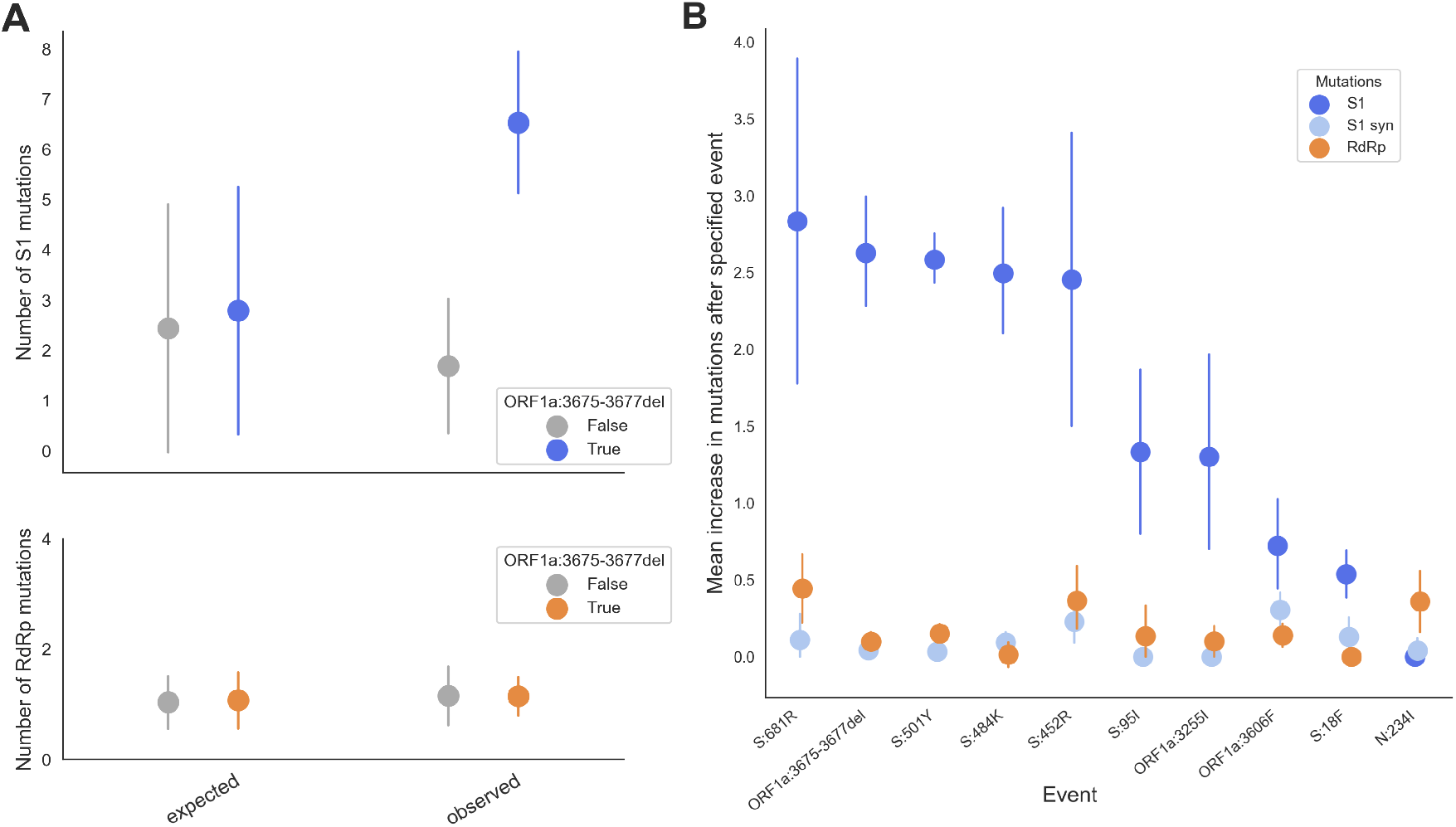
Clades with the 3-amino acid deletion in Nsp6 have a high number of S1 mutations. **A)** The mean number of S1 mutations (top), or RdRp mutations (bottom), that occur in clades that have (blue/orange) or do not have (gray) the 3-amino acid deletion in Nsp6. The expected difference is shown on the left, and empirical data is shown on the right. Expectation is based off of 100 randomizations of the tree. Error bars shown standard deviation. **B)** The difference in the number of nonsynonymous S1 (dark blue), S1 synonymous (light blue), and nonsynonymous RdRp (orange) mutations that occur before versus after a convergently-evolved mutation is shown. Error bars show 95% confidence intervals.

That clades with ORF1a:3675-3677del have higher numbers of S1 substitutions does not speak to the directionality of this relationship. In other words, it is possible that ORF1a:3675-3677del occurs in lineages that already have a lot of S1 substitutions, or that a lot of S1 mutations accumulate in clades that already have ORF1a:3675-3677del. To determine the directionality of this difference, we considered every phylogenetic path that contains the Nsp6 deletion and found the difference between the final number of S1 substitutions on that path and the number of S1 substitutions that had accumulated before the deletion. On average, around 2.5 S1 nonsynonymous mutations accumulate after ORF1a:3675-3677del (Figure 5B). This is the second largest increase in S1 mutation accumulation following any convergently-evolved mutation, behind S:681R. These results do not indicate that the deletion directly causes S1 substitutions, but they do add to the observations of convergent evolution and high clade growth rates in suggesting that ORF1a:3675-3677del is an adaptive mutation and an influential factor in the evolution of SARS-CoV-2.

## Discussion

Detecting adaptive evolution is both highly interesting from a basic scientific perspective as we seek to understand how and when this type of evolution occurs, and highly relevant from a public health perspective as we strive to curb the transmission of infectious diseases. As widespread SARS-CoV-2 circulation continues, our best defense is through vaccination. The SARS-CoV-2 vaccines showed high efficacy in clinical trials, but we must be proactive to ensure their continued effectiveness. Vaccines against viruses that undergo antigenic drift, like influenza, must be continually updated to match circulating variants. Therefore, the propensity of SARS-CoV-2 to evolve adaptively in spike S1 (the location of most neutralizing antibody epitopes) has important bearing as to whether the SARS-CoV-2 vaccine will also need to be regularly updated.

SARS-CoV-2 exhibits convergent evolution [9–11], and some of the notable mutations that have occurred multiple times independently (like S:501Y and S:484K) appear in multiple VOCs, suggesting positive selection on these mutations. In the context of deep mutational scanning (DMS) experiments, mutations at 501 increase ACE2 binding affinity [14] and mutation to site 484 escapes antibody binding [15]. Recurrent mutations at S:681 enhance S1/S2 subunit cleavage [16, 17], a protein-modification that is essential for spike-mediated cell entry [18] and thus is thought to contribute to increased viral replication [17]. Many other convergently-evolved mutations are also shared by VOCs and possess demonstrably different phenotypes, often altering antigenicity [19–21].

Despite the demonstrably advantageous effects of observed mutations, it is too soon, evolutionarily, to pick up strong signals of adaptive evolution by classical methods that rely on fixation of nonsynonymous mutations. Instead, we capitalize on the high temporal and geographic density of SARS-CoV-2 sequencing data to create a new method for identifying adaptive evolution and regions of the genome where this evolution is localized. This method identifies genes where amino acid substitutions significantly correlate with clade growth rate. This can be intuitively interpreted as genes with high rates of amino acid substitutions (suggestive of positive selection) that result in more successful viruses (suggestive of a positive fitness effect) are undergoing adaptive evolution and is effectively a continuous analog to partitioning differences between polymorphism and divergence in the classical McDonald-Kreitman test [22]. We find that the spike S1 subunit shows strong signals of adaptive evolution by this method (Figure 1).

Our inference of adaptive evolution is based on a correlation between S1 substitution accumulation and clade success that falls well outside the null expectation (Figure 1C). It is important to emphasize that these results speak to the average evolutionary effect of S1 substitutions. This does not mean that every S1 substitution is selectively advantageous, and it is likely that some mutations have larger effects on fitness than others. In fact, it is possible that successful viruses contain some S1 substitutions that do not contribute to their evolutionary success. One possibility is that these mutations could have arisen during long-term infections where they were advantageous within a single host. For instance, S1 mutations 484K and 501Y have been observed to arise from continued evolution within a single host [23]. It is therefore possible that the parallel evolution of these particular mutations is due to a selective advantage at a within-host, rather than between-host, level. Within-host selection pressures may help to explain why some mutations such as 484K occur again and again across the phylogeny (Figure 4). However, the context in which S1 mutations arose does not affect our finding that viruses with more nonsynonymous mutations in S1 are more successful, on average, within the global population of SARS-CoV-2 viruses.

Phylogenetic inferences of evolution can be biased by the samples included in the analysis. To reduce sampling biases, our study is based on a phylogeny of 9544 SARS-CoV-2 genomes sampled evenly over space and time. The strong correlation between S1 mutation accumulation and clade growth rate persists if the number of genomes included in the phylogeny is doubled (Figure S10), indicating that our results are not biased by the number of samples included in the analysis. We also find that global adaptive evolution in S1 is not driven solely by certain geographic regions. Using phylogenies that only include samples from a particular geographic region, we observe that clade success strongly correlates with S1 substitutions in Asia, Europe, North America, Oceania and South America (Figure S11). The only region where this correlation is not observed at this timepoint is Africa, where decline in frequency of a particular clade of Beta drives an overall lack of correlation.

In addition to sampling biases, there are several limitations to our approach presented here. Firstly, our analysis intentionally considers the average effect of mutations in different regions of the genome on viral fitness, with the goal of taking a populations genetics approach to quantify adaptive evolution of SARS-CoV-2. This means that, while we observe a significant correlation, given a correlation coefficient of *r* = 0.46, our results cannot predict the fitness of specific variants solely based on S1 mutation counts. Similarly, as mentioned above, this means it is likely that some successful viral clades contain S1 substitutions that are not advantageous, but rather, are hitchhiking along with positively-selected mutations. Additionally, our analysis focuses on the period of VOC and VOI emergence from December 2019 to May 2021. So, while we can speculate how our findings of high adaptive potential in S1 will translate to future evolution of the virus, we cannot directly predict how the pace of adaptive evolution will change over time.

We observe temporal structure in the adaptive evolution of SARS-CoV-2. We find that the correlation between clade success and S1 substitutions changes over time, though shows strong signals of adaptive evolution from Jan to Sep 2021 (Figure S12). Enrichment of the ratio of nonsynonymous to synonymous divergence (*d*_*N*_*/d*_*S*_) in S1 also increases over time (Figure 2). Additionally, substitutions within S1 cluster temporally (Figure 3), rather than accruing at a steady rate. This temporal structure potentially indicates a changing evolutionary landscape: either through the emergence of new selective pressure, and/or through the occurrence of permissive mutations that made adaptive mutations more accessible.

While the overall *d*_*N*_*/d*_*S*_ ratio in S1 is 0.70, *d*_*N*_*/d*_*S*_ is 1.66 along persistent lineages in 2021 (Figure 2). This high ratio is remarkable when compared to the antigenically-evolving HA1 subunits of influenza H3N2 and H1N1pdm or the S1 subunit of seasonal coronavirus OC43 (Figure S5). We estimate the mean *d*_*N*_*/d*_*S*_ ratio for HA1 in influenza H3N2 to be 0.36 (Figure S5), which is similar to the 0.37 estimated previously [8]. However, influenza H3N2 has been endemic in the human population for over 50 years, and its current evolution is largely driven by antigenic changes [24].

Viral evolution directly after spillover into a new host species can differ from evolution at endemicity. For instance, for 2 years following its emergence in 2009, the evolution of influenza H1N1pdm is thought to largely have been adaptation to a new host, whereas adaptive evolution after 2011 has been dominated by antigenic changes [25]. Accordingly, we observe that *d*_*N*_*/d*_*S*_ in the H1N1pdm HA1 subunit peaks at 0.72 in 2010 shortly after pandemic emergence, and then declines to a more stable value of ∼0.3 beginning in 2014. An initially high rate of protein-coding changes is consistent with the idea that, soon after a spillover event, there are many evolutionarily-accessible mutations that are advantageous in the new host environment. It is unclear whether the observed *d*_*N*_*/d*_*S*_ ratio in SARS-CoV-2 S1 will persist or whether it is a feature of this virus’s recent emergence and will drop in the years to come.

Together, the results presented in Figures 1-3 offer phylogenetic evidence that SARS-CoV-2 is evolving adaptively and that the primary locus of this adaptation is in S1. Our results are consistent with experimental demonstration of phenotypic changes conferred by VOC spike mutations [15, 17, 19, 21]. Adaptive evolution in the S1 subunit during the period we focus on (December 2019 to March 2021) is likely driven by selection to adapt to a new host by increasing infectivity of human cells. However, the amount of immunity to SARS-CoV-2 is rising globally, increasing the selection for antibody escape. Given the virus’s demonstrated propensity for adaptive change in S1, antigenic drift will likely begin to sculpt the evolution of SARS-CoV-2. The potential antigenic impact of adaptive S1 mutations, which are accruing at pace over 4 times that of influenza H3N2 (Figure 2, Figure S5), suggests that it may become necessary to update the SARS-CoV-2 vaccine strain. Indeed, the emergence of the Omicron Variant of Concern demonstrated a SARS-CoV-2 virus with an extraordinarily high number of S1 substitutions [26] that spread rapidly across the world and showed significantly reduced neutralization titers relative to preceding variants [27]. With Omicron, we now know that significant antigenic variants can emerge with highly modified S1 domains. However, the observed pace of adaptive evolution in S1 perhaps should have suggested the potential for emergence of such a variant.

In addition to S1, our results suggest that substitutions within Nsp6 and ORF7a may significantly contribute to the success of viral clades (Table 1). We expand on these gene-wide results by identifying specific adaptive mutations, using the confluence of convergent evolution and clade success. This analysis turned up many S1 mutations that have been extensively studied, along with mutations to nucleocapsid (N), another target of antibody-recognition [28], and a couple mutations in Nsp6, Nsp4 and M (Figure 4). The nonS1 mutations ORF1a:3255I (in Nsp4), M:82T, and N:205I in particular show compelling evidence of positive selection. These sites enriches our understanding from gene-wide analyses presented in Figures 1-3 and Table 1: though S1 is the primary genomic locus of adaptive evolution, a handful of positively-selected mutations in other genes are also influencing the evolution of SARS-CoV-2 in the human population.

Our analysis of specific adaptive mutations suggests the possibility of differences between within-host selection for viral replication and between-host selection for transmission. Viruses belonging to Delta have shown greater between-host transmission rates than other VOC or VOI viruses [29], but are lacking mutations that have occurred repeatedly and that were associated with increased clade growth (notably ORF1a:3675-3677del, S:484K and S:501Y). It is possible that some mutations display a large degree of parallelism due to specific within-host pressures that occur in secondary infections of partially immune individuals, despite having only modest effects on between-host transmission.

It is important to note that the precise mutations that appear most influential depend on when the analysis is done (Figures 4C and S9). The fitness effect of a mutation is not an absolute quality — it depends on a multitude of influences including genetic background of the viral lineage, other co-circulating lineages, existing host immunity, and epidemiological factors (such as geographically heterogeneous mitigation efforts). Additionally, lineages can grow in frequency due to stochastic effects. It is, therefore, expected that mutations associated with successful clades will change over time and that these changes reflect both a changing fitness landscape and the stochastic nature of evolution. Mutations that transcend this or, in other words, are associated with successful lineages at multiple time points, are more likely to have important, adaptive functions. One such mutation is ORF1a:3675-3677del (Figures 4C and S9).

The ORF1a:3675-3677 deletion removes 3 amino acids (SGF) from a predicted transmembrane loop [30] of the Nsp6 protein. Across the coronavirus family, the Nsp6 protein, in coordination with Nsp3 and Nsp4, forms double-membrane vesicles that are sites for viral RNA synthesis [31]. In SARS-CoV-2, Nsp6 suppresses the interferon-I response [32]. It is unclear whether ORF1a:3675-3677del impacts either of these functions. This deletion is not observed in other sarbecoviruses, residues 3675 and 3676 are 100% conserved, and only synonymous and conservative changes are seen at 3677 in this subgenus [33]. However, in SARS-CoV-2, this deletion exhibits close to the highest level of convergence, presence in VOCs, mean logistic growth rate, and increase in S1 mutations in descending lineages. Future experimental study of this deletion would increase our understanding of what functions, apart from enhanced cell entry and potential antibody escape, were highly advantageous during the early adaptive evolution of SARS-CoV-2.

So far, ORF1a:3675-3677del has not been observed in Delta viruses and our results suggest that the appearance of a sublineage of Delta possessing ORF1a:3675-3677del may outcompete basal Delta viruses. However, the Omicron variant, which appeared in late November 2021 and spread rapidly, possesses a very similar deletion, where Omicron viruses from the primary BA.1 PANGO lineage exhibit ORF1a:3674-3676del. This provides yet another example of deletion to this region of Nsp6 being associated with consequential VOC viruses.

## Methods

Source code for all analyses presented in this manuscript is available at github.com/blab/sarscov2-adaptive-evolution.

### Phylogenetic reconstruction of a subsampling of global SARS-CoV-2 genome sequences

All primary analyses in this manuscript were performed using data downloaded from the GISAID EpiCoV database (gisaid.org, [2]) on July 29, 2021 and curated by the Nextstrain nCoV ingest pipeline (github.com/nextstrain/ncov-ingest). This dataset contained 2,459,376 viral genomes and associated metadata. These genomes were aligned with Nextalign (docs.nextstrain.org/projects/nextclade/en/latest/user/nextalign-cli.html) and masked to minimize error in phylogenetic inference associated with problematic amplicon sites. Masked alignments were filtered to exclude strains that were known outliers, sequenced due to ‘S dropout’, mis-annotated with an admin division of ‘USA’, shorter than 27,000 bp of A, C, T, or G bases, missing complete date information, annotated with a date prior to October 2019, flagged with more than 20 mutations above the expected number based on the mutational clock rate, or flagged by Nextclade (docs.nextstrain.org/projects/nextclade/en/latest/user/algorithm/07-qualitycontrol.html) with one or more clusters of 6 or more private differences in a 100-nucleotide window. After filtering 2,213,085 genomes remained.

After filtering, SARS-CoV-2 genomes were evenly sampled across geographic scales and time. Specifically, a maximum of 1,600 strains were sampled from each continental region including Africa, Asia, Europe, North America, Oceania, and South America for an approximate total of 9,600 genomes per phylogeny. For each region except North America and Oceania, strains were sampled from each distinct combination of country, year, and month. For North America and Oceania, genomes were sampled from each distinct combination of division (i.e., state-level geography), year, and month.

Time-resolved phylogenies were inferred using Augur 12.0.0 [34], IQ-TREE 2.1.2 [35], and TreeTime 0.8.2 [36]. Ancestral sequences were inferred with TreeTime using the joint inference mode. The primary analysis was conducted on 9544 genomes collected on or before May 15, 2021, and the phylogeny reconstructed from these data can be found at nextstrain.org/groups/blab/ncov/adaptive-evolution/2021-05-15. Phylogenies used for secondary analyses of convergent evolution (Figure 4C, and Figure S9) can be viewed using the date drop-down menu in the left-hand sidebar. The secondary analyses included isolates sequenced up until April 15, 2021 (9467 genomes), May 1, 2021 (9449 genomes), June 1, 2021 (9343 genomes), and June 15, 2021 (9401 genomes). All isolates used in these analyses are listed in the Acknowledgements table in the Supplementary Material.

### Quantification of mutation accumulation

For every internal branch on the phylogeny, the number of mutations that accumulated between the root of the tree and that branch was counted. For this and all subsequent analyses, deletions are grouped with nonsynonymous substitutions. Deletions that span multiple, adjacent amino acids are counted as one mutation. Mutations to a premature stop codon are also counted as one mutation event. Mutations were separated by which gene they occur in (according to the Wuhan-Hu-1 reference sequence, found at `analysis/reference_seq_edited.gb`) and whether they are synonymous or nonsynonymous. Genomic locations of the 15 NSPs were found in the NC 045512.2 annotation of the ORF1ab polyprotein (www.ncbi.nlm.nih.gov/gene/43740578). Code for mutation accumulation counting and plotting of Figure 1A is found in fig1-muts_by_time_and_growthrate.

### Estimation of the logistic growth rate of clades

Logistic growth of individual clades was estimated from the time-resolved phylogeny and the estimated frequencies for each strain in the tree. Frequencies were estimated with Augur 12.0.0 [34] using the KDE estimation method that creates a Gaussian distribution for each strain with a mean equal to the strain’s collection date and a variance of 0.05 years. At weekly intervals, the frequencies of each strain at a given date were calculated by summing the corresponding values in their Gaussian distributions and normalizing the values to sum to 1. The frequency of each clade at a given time was the sum of its corresponding strain frequencies at that time.

Logistic growth was calculated for each clade in the phylogeny that was currently circulating at a frequency *>*0.0001% and *<*95% and that had at least 50 descendant strains. Each clade’s frequencies for the last six weeks were logit transformed and used as the dependent variable for a linear regression where the independent variable was the corresponding date value for each transformed frequency. The logistic growth of the clade was then annotated as the slope of the linear regression of the logit-transformed frequencies.

### Calculation of nonsynonymous to synonymous divergence ratio

A time-course of *d*_*N*_*/d*_*S*_ ratios was calculated in non-overlapping time windows by splitting all internal branches included in the phylogeny according to their date. Within each gene, the nonsynonymous and synonymous Hamming distances were found between the reference sequence and every internal branch. The Hamming distances were normalized by the total number of possible nonsynonymous or synonymous sites within that gene to give a measure of divergence. The nonsynonymous divergence was divided by synonymous divergence. Then, for each time window, the mean of this ratio was found for all internal branches within the window. For SARS-CoV-2, the time windows were 2 months and overlap by 1 month. The code to run this analysis and reproduce Figure 2 is at `fig2-divergence.ipynb`.

Phylogenies for seasonal influenza A/H3N2 and A/H1N1pdm were built using the Nextstrain pipeline from github.com/nextstrain/seasonal-flu. They include 2274 and 2169 genomes, respectively, that were sampled between 2009 and 2021 to capture the earliest sequences from the H1N1pdm pandemic (March 2009). The OC43 phylogeny was built from all available OC43 lineage A genomes sampled in 2009 or later (214 genomes) using the workflow in github.com/blab/seasonal-cov-adaptive-evolution/oc43/separate lineages. Divergence accumulation ratios were computed from the root of each tree using 1-year time windows overlapping by 0.5 years. These phylogenies can be found in `seasonal-flu_trees/`. The code in `fig2supp-divergence_seasonalflu.ipynb` reproduces Figure S5.

### Randomization of mutations across the phylogeny for wait time calculations

For each type of mutation (S1 nonsynonymous, S1 synonymous, and RdRp nonsynonymous), the total number of mutations observed on the phylogeny was randomly scattered across phylogeny. Only internal branches with 3 or more descending tips were used. Random branches were selected by a multinomial draw, where the likelihood of a branch having a mutation is proportional to its branch length in years. Multiple mutations were allowed to occur on the same branch, just as with the empirical phylogeny. Randomizations were run 1000 times for each mutation type used in Figure 3B and C, and 10 times for the distributions shown in figure S7. Code for this analysis is in `fig3-wait_times.ipynb`.

### Calculation of wait times

Wait times were counted for the following classes of mutations: S1 nonsynonymous, S1 synonymous, and RdRp nonsynonymous. For each class of mutation, a wait time was calculated between each branch that has a mutation of this type and its first child branch on each descending path that has a mutation of this type. A wait time was also calculated between the tree root and the first branch on any independent path that has a mutation of this type. Conceptually, the result of this is that wait times are computed between every sequential mutation that occurs along every path on the tree (as diagrammed in Figure 3A), without double counting any pairs of branches. Only mutations on internal branches (defined as having 3 or more descending tips) are considered.

A wait time is simply the time between mutations and is calculated by subtracting the date (in decimal years) of the earlier mutation from the date of the later mutation. Because the exact date a mutation occurred cannot be known, each mutation is assigned a random date along the branch it occurred on. If multiple mutations of the same type occurred on one branch, each mutation is assigned a different random date and the wait times between mutations on that branch are calculated.

Empirical and expected wait times were calculated for each type of mutation 1000 times and the results of all 1000 iterations can be found in `wait_time_stats/`. Code to calculate wait times and reproduce Figure 3B and C and Figure S7 is found in fig3-wait_times.ipynb.

### Quantification of convergent evolution and logistic growth rates across the phylogeny

Every substitution that occurred on an internal branch with at least 15 descending tips was tallied. For every substitution that was observed at least 4 times on internal branches, the average growth rate of clades containing this mutation was calculated by taking the mean logistic growth rate of clades where this mutation occurred. Code to count occurrences, calculate mean logistic growth, and determine which emerging lineages descend from recurrent mutations is found in fig4-convergent_evolution.ipynb. This code will reproduce Figures 4A, S8, and S9.

### Randomization of recurrent mutations across the phylogeny

One hundred randomized trees were created by shuffling the phylogenetic positions of each substitution that was observed on an internal branch with at least 15 descending tips (those calculated above and shown in Figure 4A). Randomized branches were also limited to internal branches with at least 15 descending tips. The position of each randomized substitution was constrained to branches that “make phylogenetic sense”: meaning, a given substitution cannot occur twice on the same path. This results in a tree with exactly the same distribution of mutation occurrences as the empirical phylogeny, but where those mutations occur on different branches. Code to implement these randomizations and reproduce Figure 4B is in fig4-convergent_evolution.ipynb.

### Consideration of recombination as an alternative to convergent evolution of nsp6 deletion

For each occurrence of the ORF1a:3675-3677 deletion, all nucleotide mutations that occurred between the root and the branch where the deletion occurred were recorded. Then, recombination between every pair of the 8 inferred occurrences of ORF1a:3675-3677del was considered. For each pair, informative mutations that did not occur in a common ancestor of the potential recombinant lineages were identified. The informative mutations closest to the Nsp6 deletion on the upstream side were compared between potential donor and acceptor (and the same was done for the downstream side). If the closest mutations were shared between any donor/acceptor pair, this would be evidence that this mutation and the Nsp6 deletion were transferred from the donor to the acceptor by recombination. If the closest mutations are not shared between the donor and acceptor, the only way the acceptor could have acquired the ORF1a:3675-3677del through recombination is if both recombination break points occurred within a genomic window defined by the closest informative mutations on either side of the Nsp6 deletion. Code for this analysis as well as a table summarizing the results is in `nsp6del_recombination.ipynb`.

### Calculation of the mean number of S1 mutations per clade

The phylogeny was divided into clades that have the ORF1a:3675-3677 deletion and those that do not, and the mean number of S1 and RdRp substitutions was computed for each category. The tree was limited to only branches occurring on or after the date of the first ORF1a:3675-3677del occurrence. The expectation was created by randomizing the locations of the 8 occurrences of ORF1a:3675-3677del as was done above in “Randomization of recurrent mutations across the phylogeny”. Code for this analysis is in fig5a-nsp6del s1mutations_correlation.ipynb.

### Calculation of S1 mutations that precede and follow specific mutation events

For each convergently-evolved mutation, every path through the phylogeny containing this mutation was considered. The total number of S1 mutations accumulated between the root and the occurrence of the convergently-evolved mutation is considered to be the number of S1 mutations before the event. The number of mutations after is the final number of S1 mutations present on the path. The before total is subtracted from the after total to give the increase in S1 mutations after the event. The mean of this increase is calculated for every path containing the convergently-evolved mutation. Code to implement this analysis is in fig5b-s1_muts_before_vs_after.ipynb.

### Consideration of sampling biases

The impact of phylogeny size on the results presented in this manuscript was tested by running the same analyses on a phylogeny containing twice as many samples. This phylogeny was built according to the same methods described in the “Phylogenetic reconstruction of a subsampling of global SARS-CoV-2 genome sequences” section above, except that a maximum of 3,300 strains were sampled from each continental region. This resulted in a tree with a total of 19,694 genomes, sampled on or before May 15,2021 and distributed roughly evenly over geography and time (nextstrain.org/groups/blab/ncov/adaptive-evolution/2021-05-15/20k). The correlation between mutation accumulation in 8 genes(or subunits) and clade success (Figure S10A)) is done in `fig1supp-20ktree.ipynb`. The *d*_*N*_*/d*_*S*_ ratio was calculated on the 19,694-tip tree (Figure S10B) as described in the “Calculation of nonsynonymous to synonymous divergence ratio” section, and this analysis is in `fig2supp-divergence-20k.ipynb`. Convergently-evolved mutations (Figure S10C) were identified on this larger tree in `fig4supp-convergent_evolution-20k`.

The impact of specific geographic regions on the results presented in Figure 1B was analyzed by computing the correlation coefficient *r* between S1 substitution accumulation and logistic growth rate for each geographic region separately. This was done by constructing a separation 10,000-sample tree for each of the 6 continental regions: Africa, Asia, Europe, North America, Oceania, and South America. Samples are from May 15, 2021 or earlier and are roughly evenly distributed over time. Each regional tree was built according to “Phylogenetic reconstruction of a subsampling of global SARS-CoV-2 genome sequences”, except that all sequences were restricted to that geographic region. The Asia-specific tree can be interactively-viewed at nextstrain.org/groups/blab/ncov/adaptive-evolution/2021-05-15/asia, and other regional trees can be accessed by substituting the region’s name at the end of the URL. Code to conduct the analysis presented in Figure S11 is in fig1-followup-regional.ipynb.

### Analysis of duration of correlation between clade success and S1 substitutions

The correlation coefficient *r* between S1 substitution accumulation and logistic growth rate was computed over time using 13 phylogenies spanning a year of time surrounding the primary analysis. Trees for this analysis were built according to the methods in “Phylogenetic reconstruction of a subsampling of global SARS-CoV-2 genome sequences” except that end date was changed to the 15th of the month, for each month between November 15, 2020 and November 15, 2021. The November 15, 2020 tree can be viewed at nextstrain.org/groups/blab/ncov/adaptive-evolution/2020-11-15, and all other dates can be accessed by changing the date at the end of the URL. Code to conduct the analysis presented in Figure S12 is in fig1-followup-timeseries.ipynb.

## Supporting information

Supplemental Acknowledgement Table

## Acknowledgements

We thank members of the Bedford Lab, Tami Lieberman and two anonymous reviewers for helpful feedback on the manuscript. We acknowledge the authors for originating and submitting laboratories of the sequences from GISAID’s EpiCoV database, on which this research is based. A full Acknowledgments table is available in the Supplementary Materials. T.B. is a Pew Biomedical Scholar. This work was supported by NIH R35 GM119774. K.E.K. was supported by the National Science Foundation Graduate Research Fellowship Program under Grant No. DGE-1762114.

## Supplementary Material

**Figure S1.**
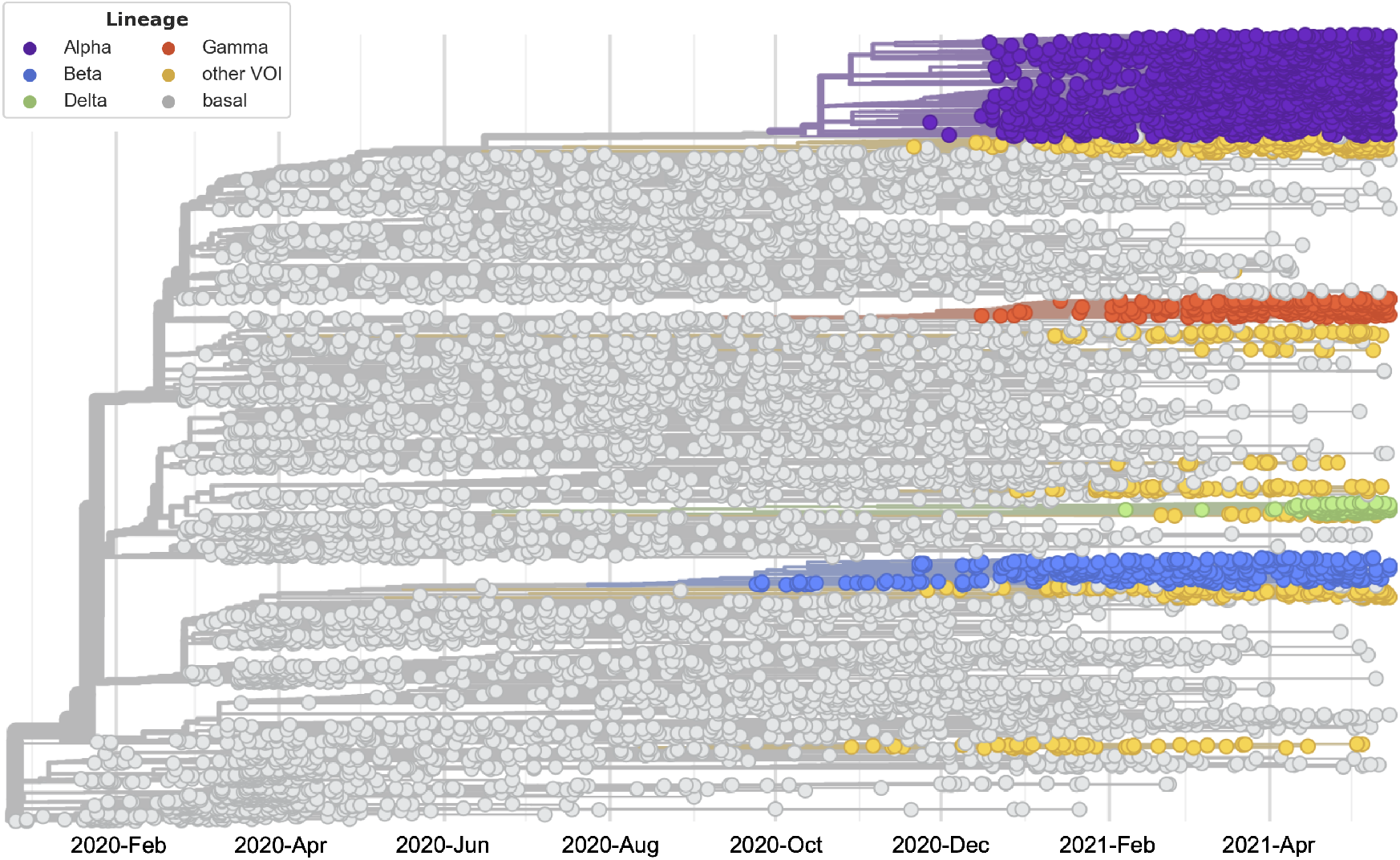
Phylogeny of 9544 SARS-CoV-2 genomes. Screenshot of the phylogeny used for the primary analyses in this manuscript. Tips and branches are colored according to viral lineage. The prominent Variant of Concern (VOC) lineages Alpha, Beta, Delta and Gamma are shown. The “other VOI” category includes 13 emerging lineages — WHO VOCS, VOIs, and prominent PANGO lineages [**?**]. Colors match those used in Figure 1. An interactive version of this phylogeny can be accessed at nextstrain.org/groups/blab/ncov/adaptive-evolution/2021-05-15.

**Figure S2.**
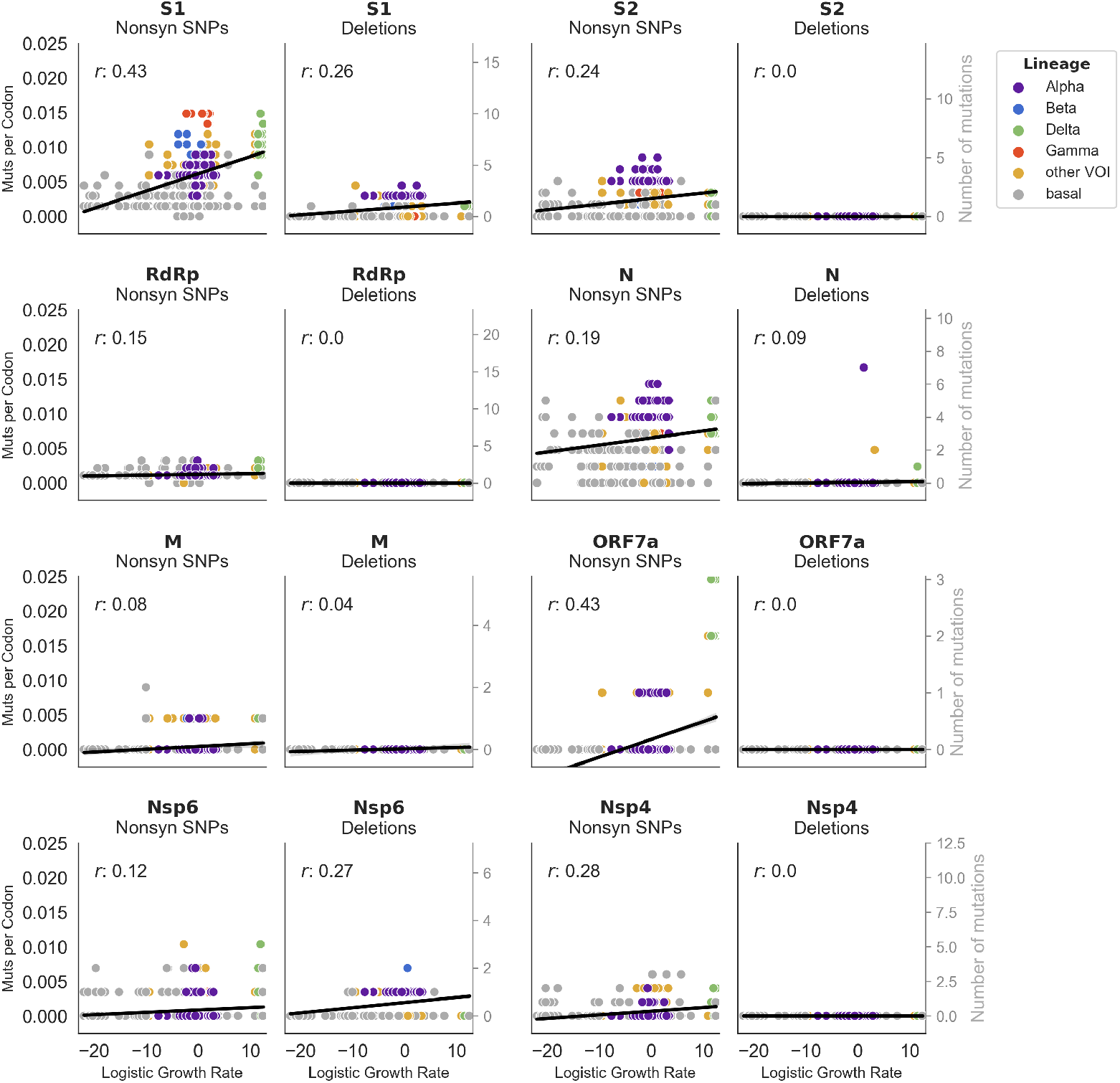
Deletions contribute to protein-coding changes in S1, N and Nsp6. For each gene nonsynonymous mutation accumulation is separated into nonsynonymous SNPs (left) and deletions (right). Accumulation of these mutations is plotted against logistic growth rate for 8 genes (or subunits), as in Figure 1B.

**Figure S3.**
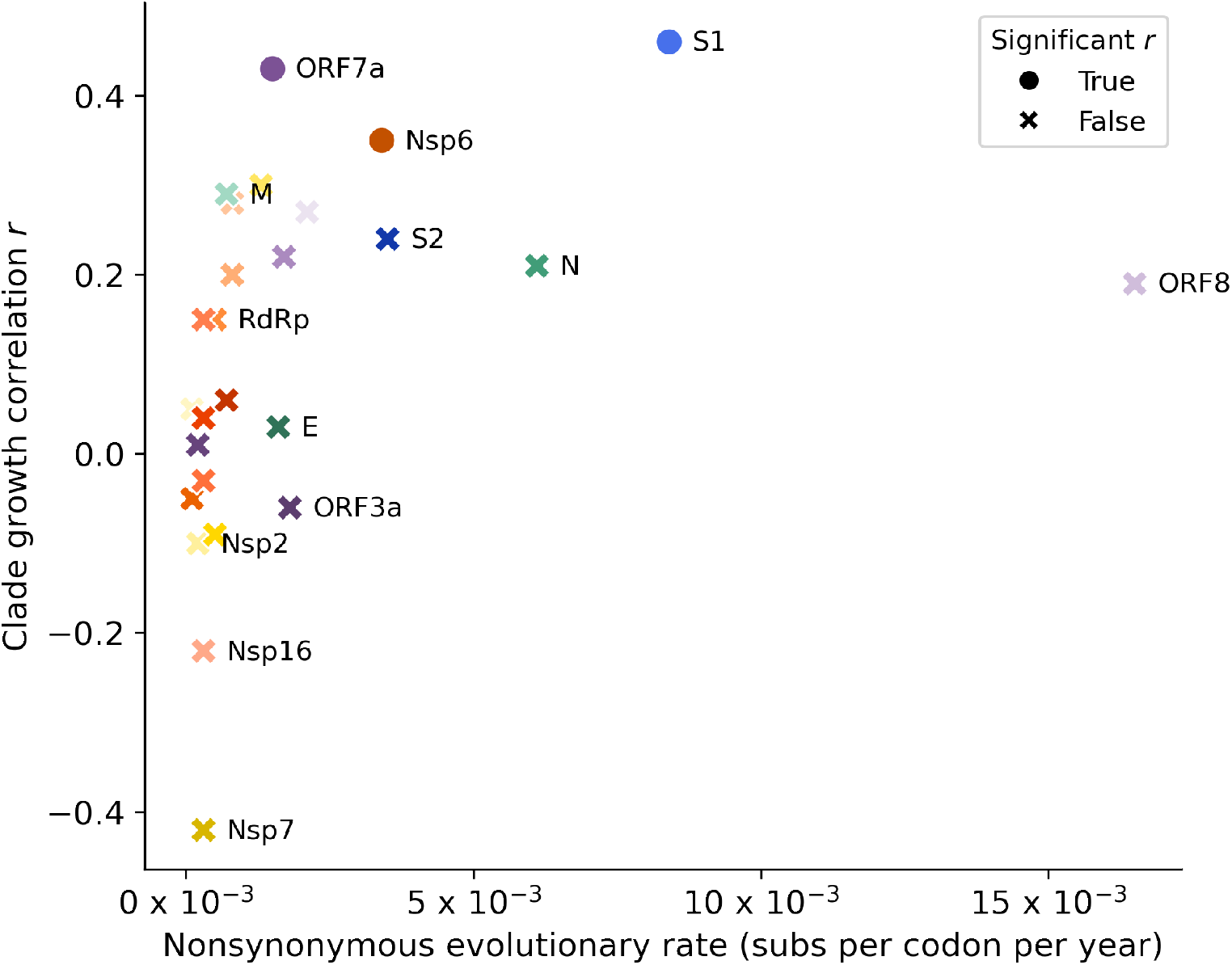
Visual Representation of Table 1. For every gene in the genome, the rate of nonsynonymous substitutions (and deletions) per codon per year is plotted against the correlation coefficient *r* of mutation accumulation with logistic growth. Circles indicate genes with significant *r* values at the *p*=0.01 level, and Xs indicate genes with insignificant *r* values.

**Figure S4.**
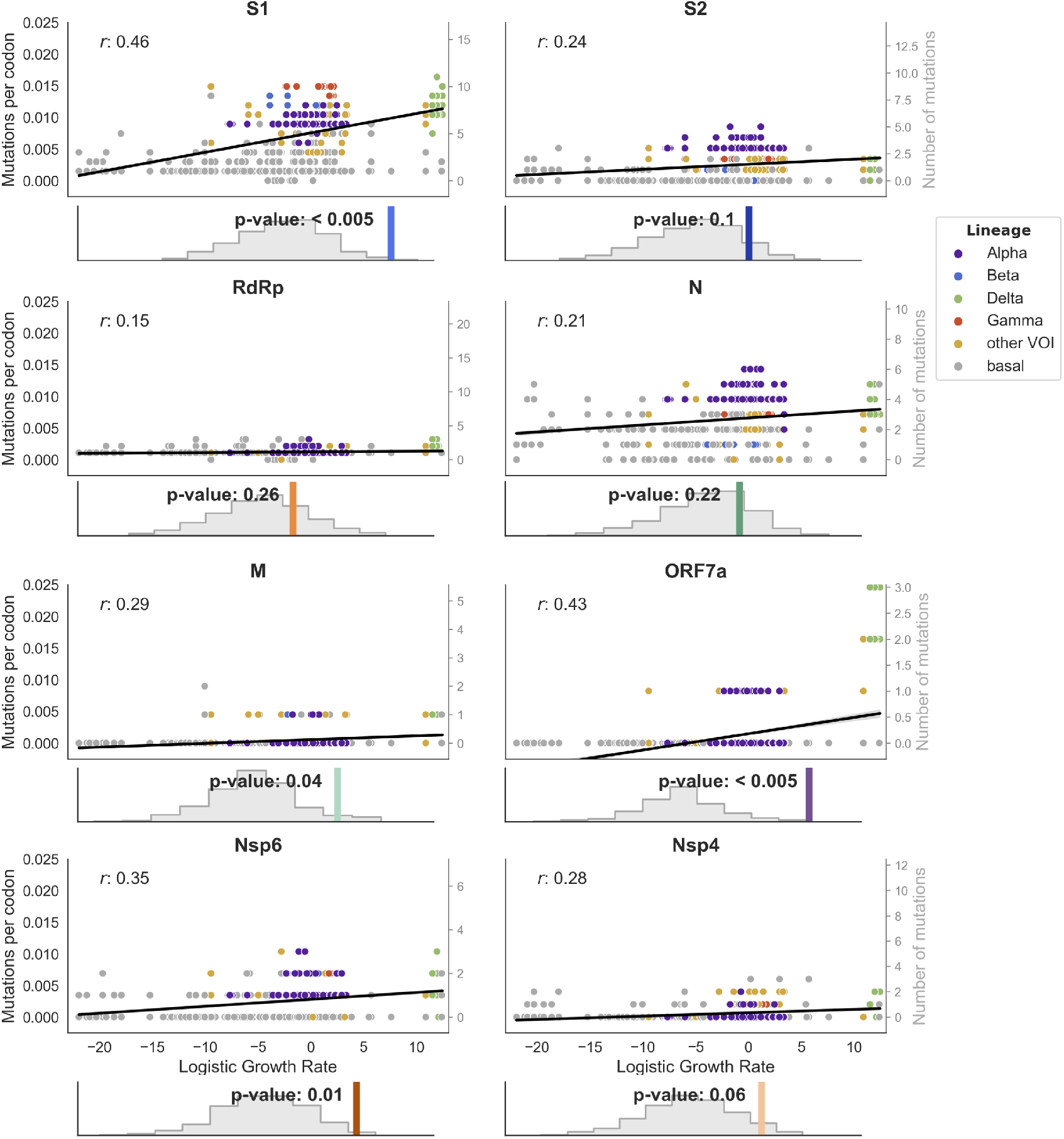
Correlation between nonsynonymous mutation accumulation and clade success is strongest in S1. Nonsynonymous mutation accumulation (mutations per codon) is plotted against logistic growth rate for 8 genes (or subunits), as in Figure 1B. Histograms beneath each plot show the empirical correlation coefficient *r* (colored line) compared to the distribution of *r* coefficients from 1000 randomizations, as well as the *p*-value resulting from this comparison.

**Figure S5.**
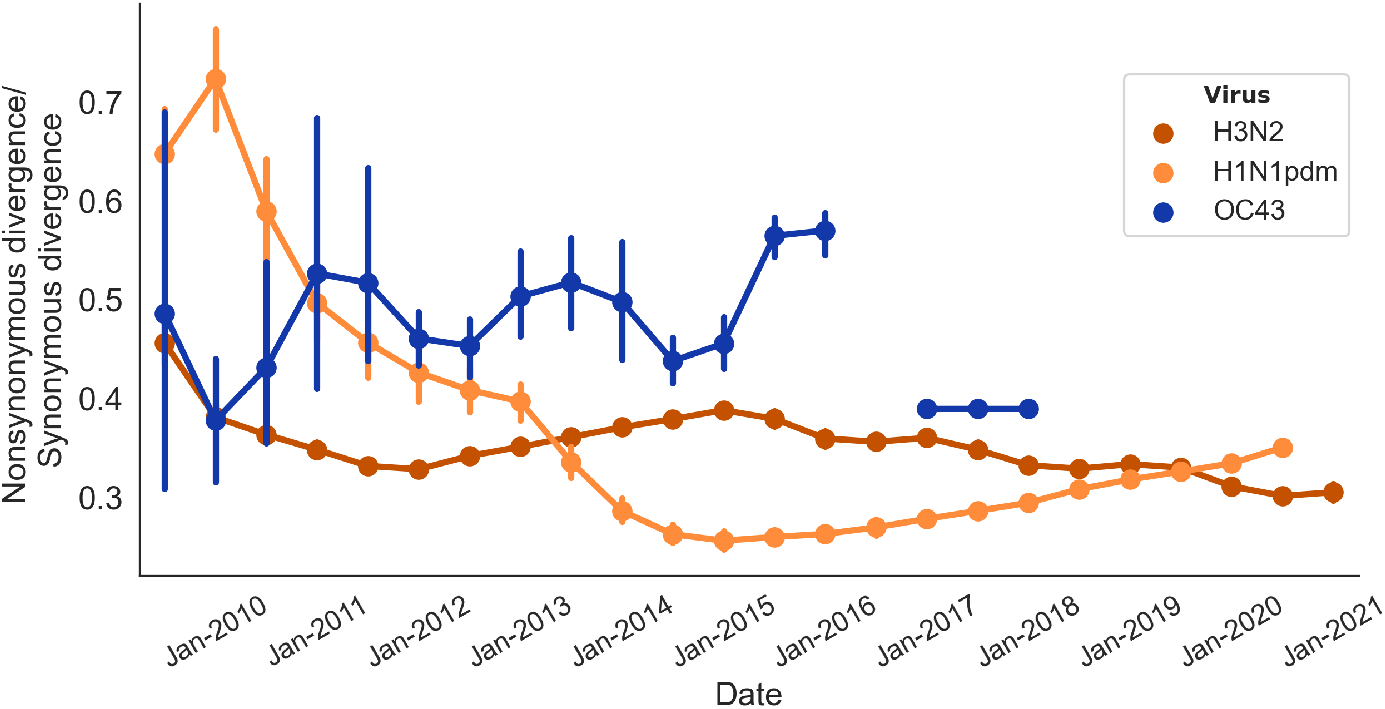
Ratio of nonsynonymous to synonymous divergence in HA1 subunit of seasonal influenza H3N2 and H1N1pdm and spike S1 subunit of seasonal coronavirus OC43. The mean and 95% confidence intervals for nonsynonymous/synonymous divergence ratios for the seasonal influenza HA1 subunits and seasonal coronavirus S1 subunit are shown over a 12-year period starting in January 2009. Divergence accumulation from the root is calculated as in Figure 2 except using windows of 1 year that overlap by half a year.

**Figure S6.**
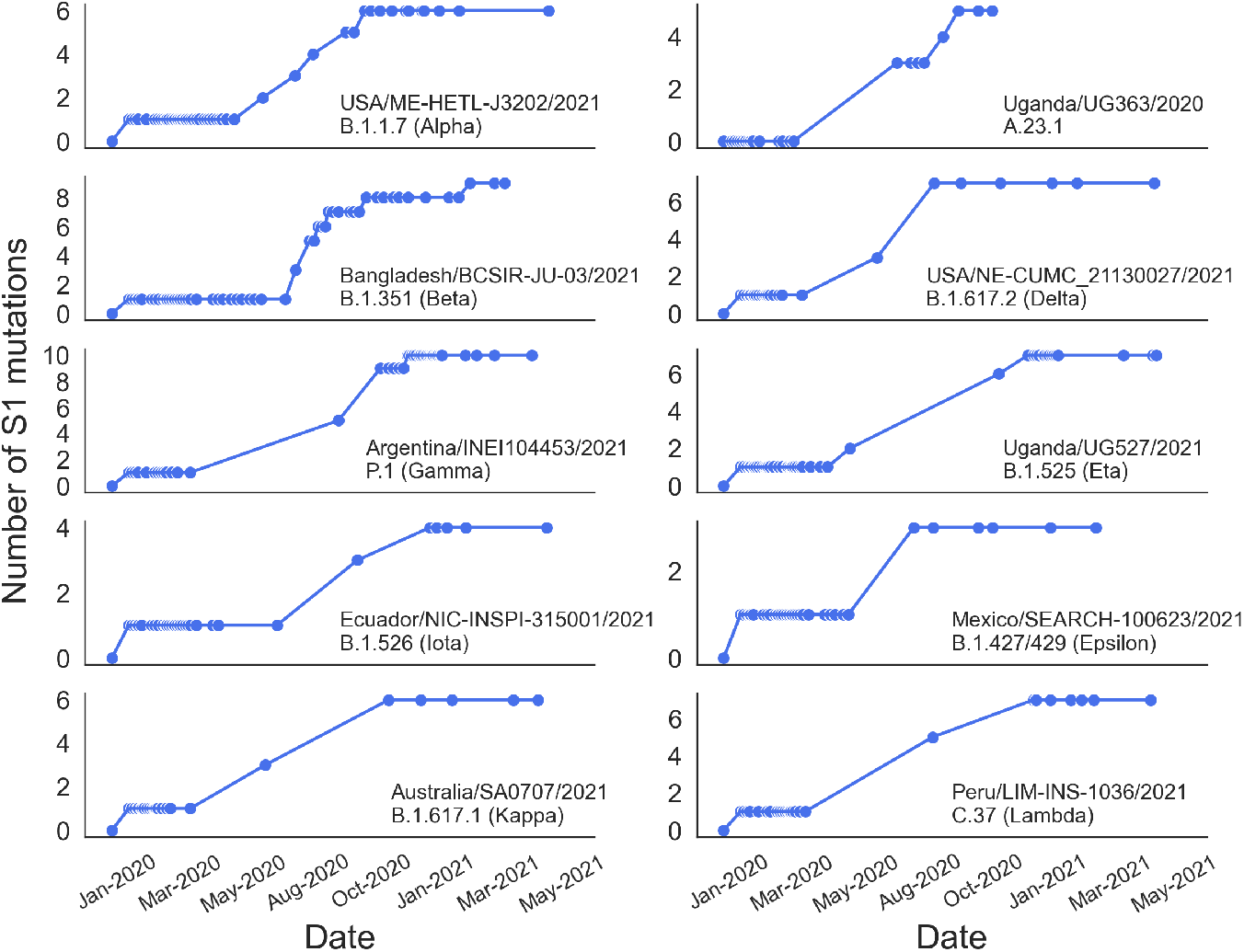
Temporal accumulation of S1 mutations on representative paths through the tree. The total number of accumulated S1 nonsynonymous mutations is counted at every branch along a path through the tree. This is plotted for 10 representative paths from the root to an isolate in an emerging lineage clade. The isolate and emerging lineage are labeled on each panel.

**Figure S7.**
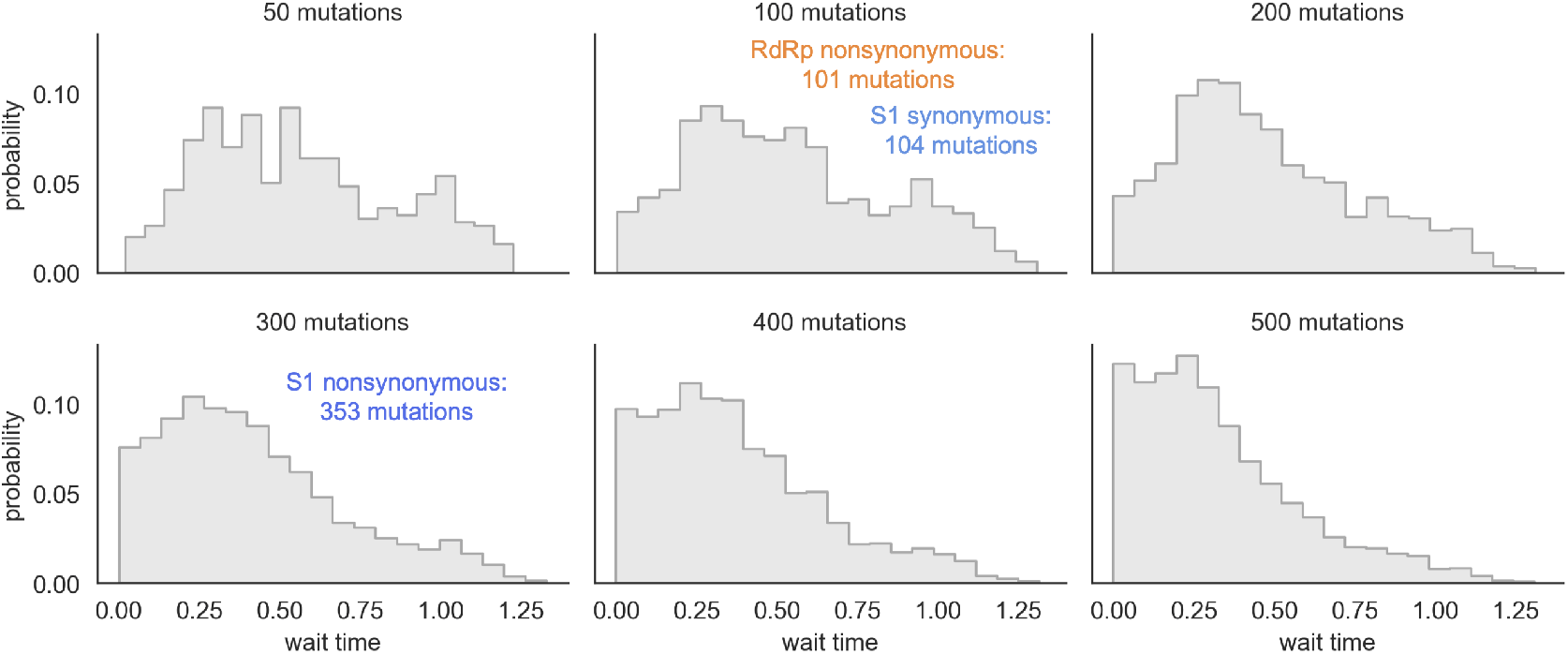
Distribution of expected wait times is affected by the number of mutations that occur across the phylogeny. The phylogeny was randomized with varying numbers of mutations to display the expected wait time distributions if 50, 100, 200, 300, 400 or 500 mutations occur on internal branches of the phylogeny. Each randomization is run for 10 iterations. The empirical number of S1 nonsynonymous, S1 synonymous, and RdRp nonsynonymous mutations observed on internal branches of the phylogeny are indicated.

**Figure S8.**
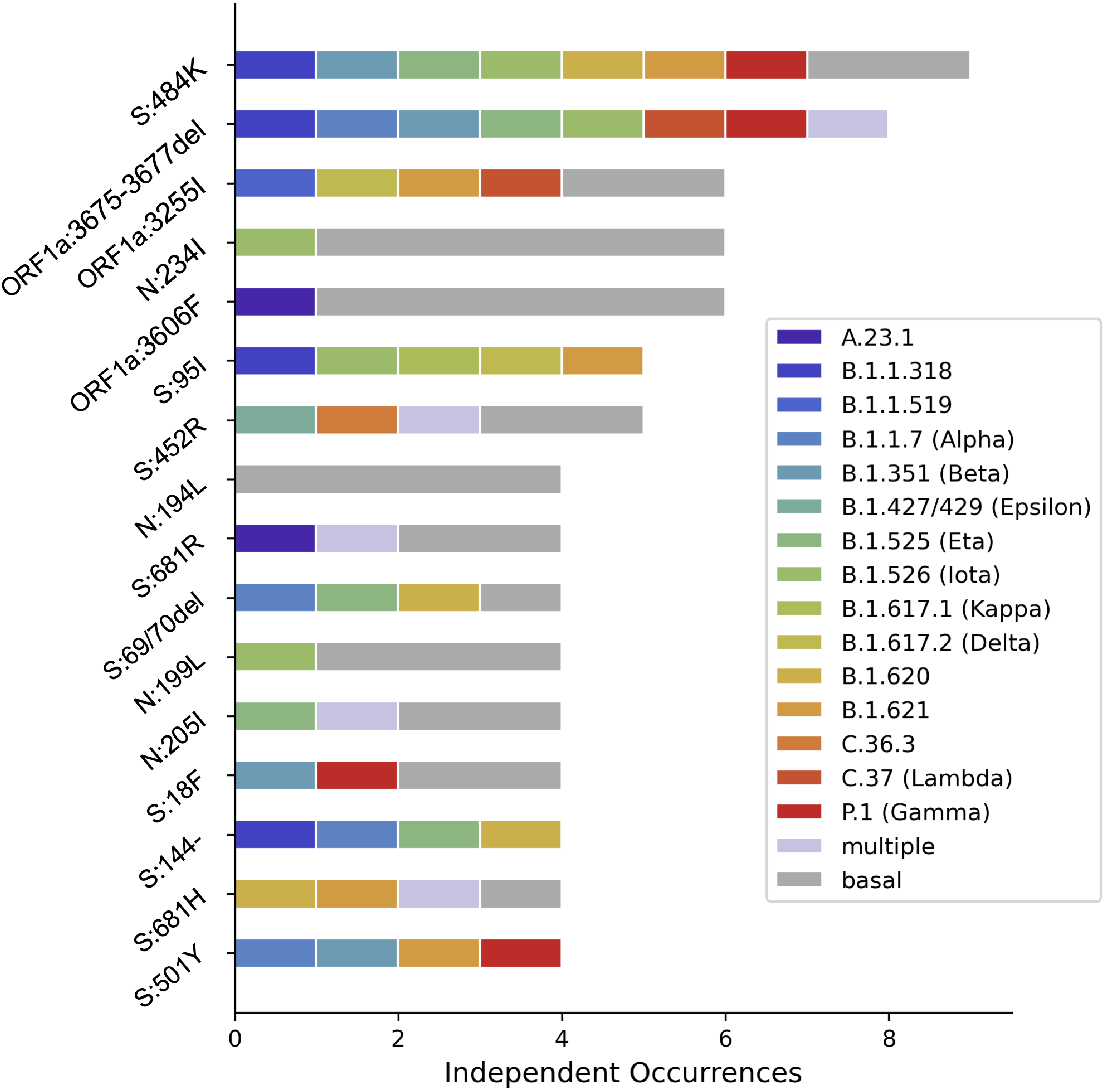
Every occurrence of the 3-amino acid deletion in Nsp6 resulted in an emerging lineage. Every occurrence of the convergently-evolved mutations is colored according to the emerging lineage it occurs at the base of. Multiple emerging lineages descending from the branch a mutation occurs on is represented by light purple.

**Figure S9.**
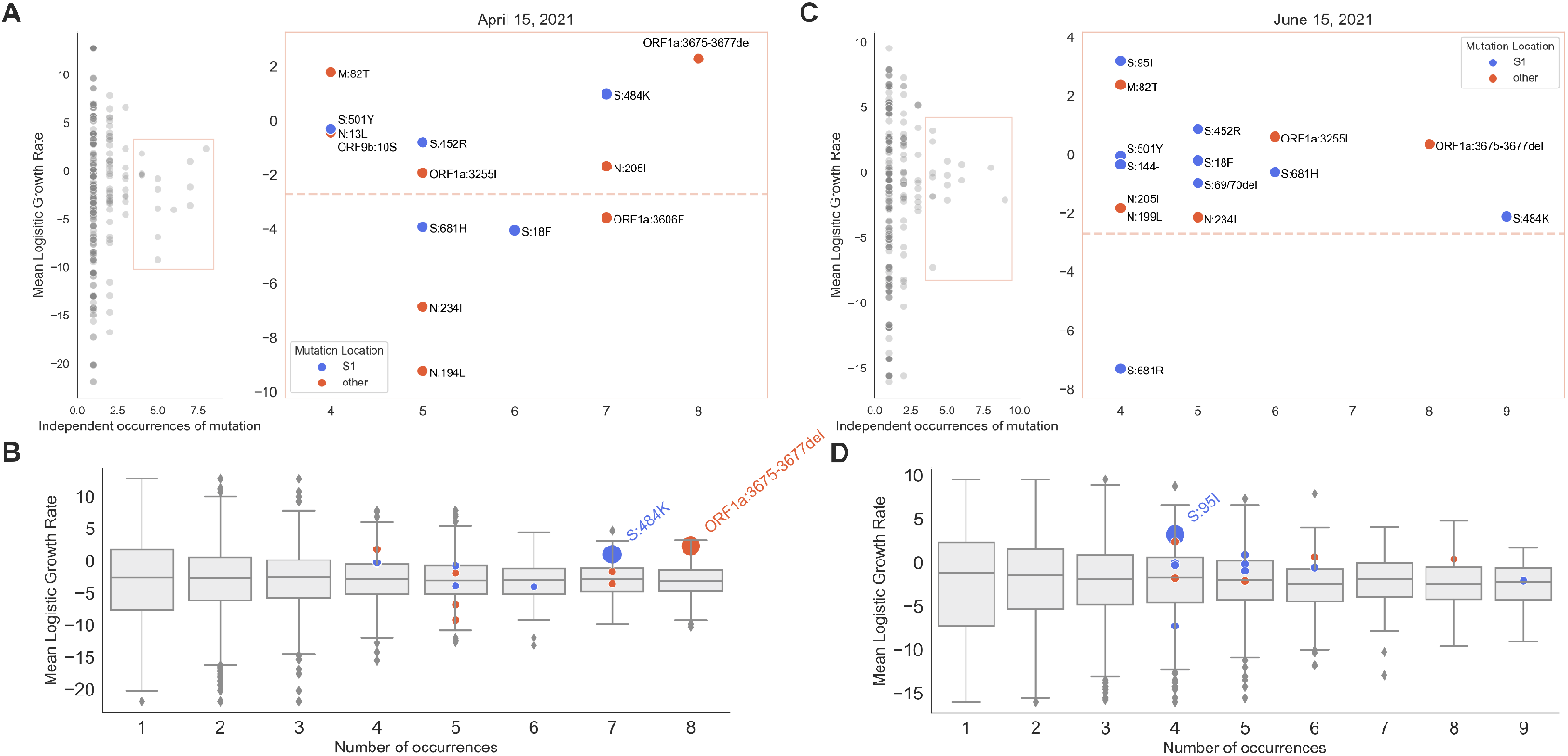
Analyses of convergent evolution shown 1 month before and 1 month after the primary analysis. **A)** Same as Figure 4A, completed using sequences up to April 15, 2021 (1 month before the primary analysis). **B)** Same as Figure 4B, completed using sequences up to April 15, 2021. **C)** Same as Figure 4A, completed using sequences up to June 15, 2021 (1 month after the primary analysis). **D)** Same as Figure 4B, completed using sequences up to June 15, 2021.

**Figure S10.**
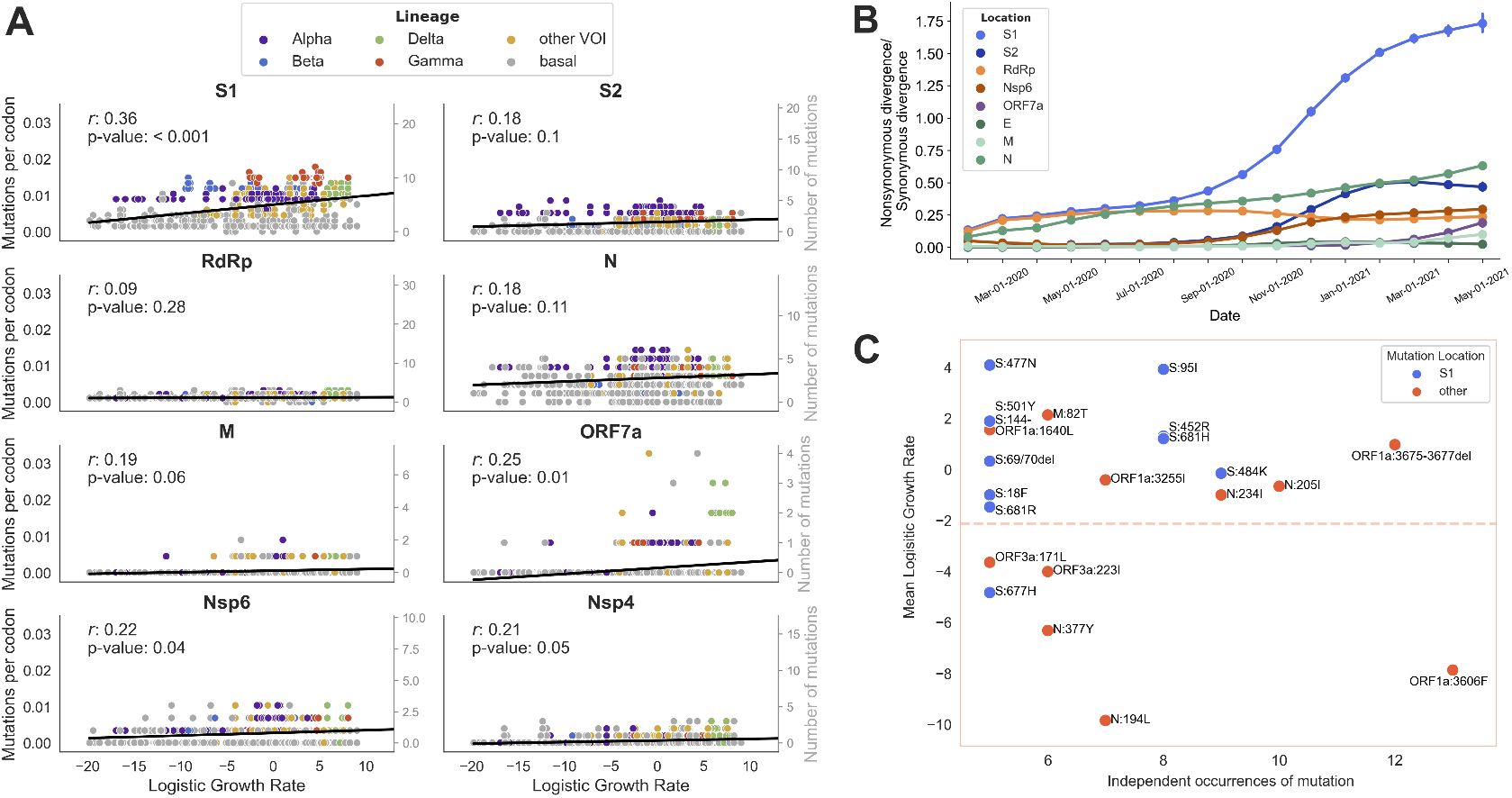
Primary results are reproduced using a phylogeny of 19,694 genomes. Analyses presented in the primary figures were repeated using a tree containing twice as many samples nextstrain.org/groups/blab/ncov/adaptive-evolution/2021-05-15/20k. **A)** Correlation between nonsynonymous mutation accumulation and clade growth for 8 genes as in Figure S4. For each gene, the empirical correlation coefficient *r* is compared to a distribution of *r* coefficients from 1000 randomizations to determine the *p*-value. **B)** Nonsynonymous to synonymous divergence accumulation ratio over time as in Figure 2. **C)** Convergently-evolved mutations that occur 5 or more times independently and the mean growth rate of every clade containing that mutation are plotted, as in Figure 4A.

**Figure S11.**
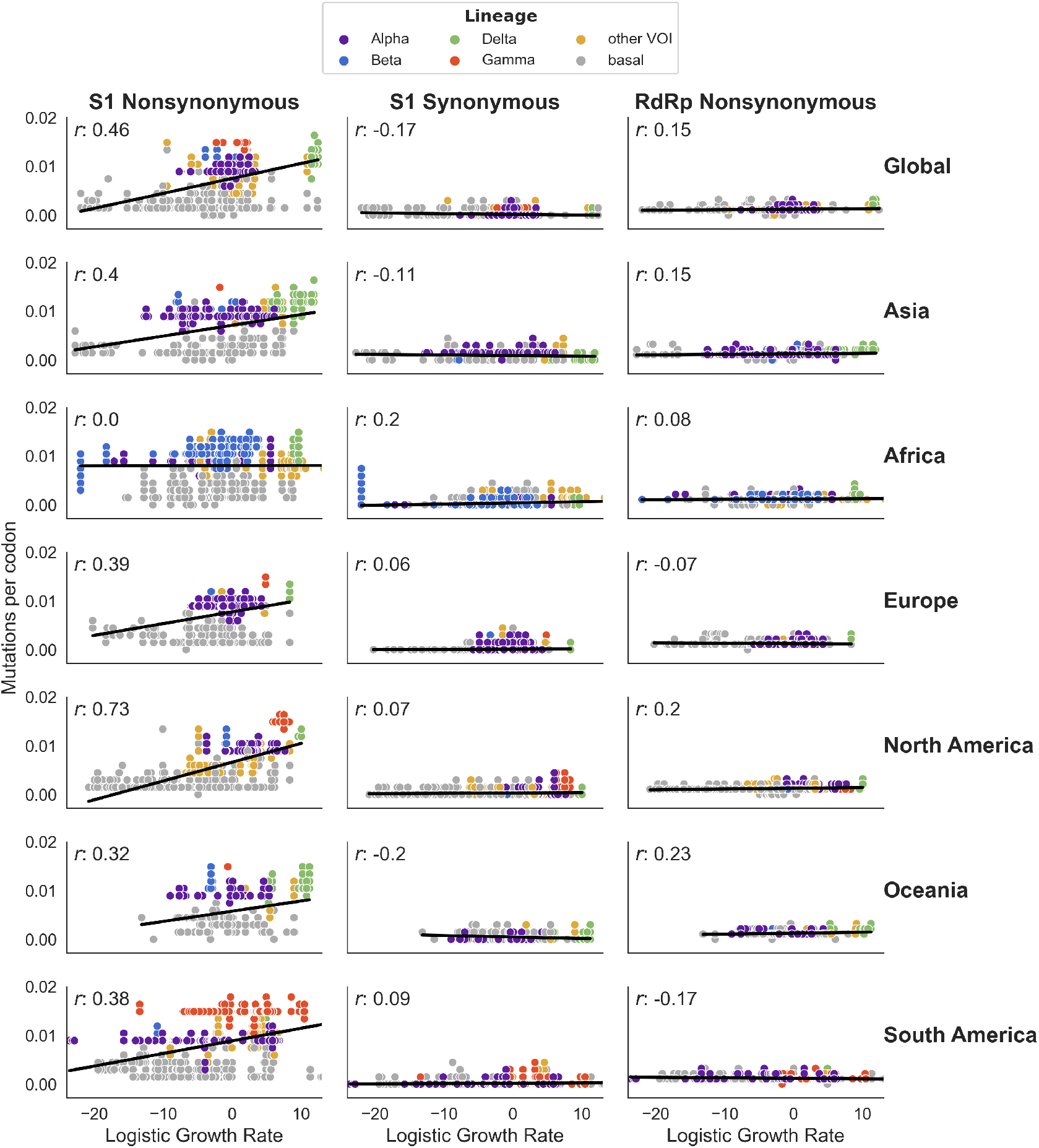
Correlation between S1 substitutions and clade growth rate is consistent across all geographic regions except one. Phylogenies were built using only around 10,000 samples from a single geographic region (Africa, Asia, Europe, North America, Oceania, and South America). The correlation between S1 nonsynonymous, S1 synonymous and RdRp nonsynonymous mutation accumulation and clade growth rate (as in Figure 1) is plotted for each geographic region.

**Figure S12.**
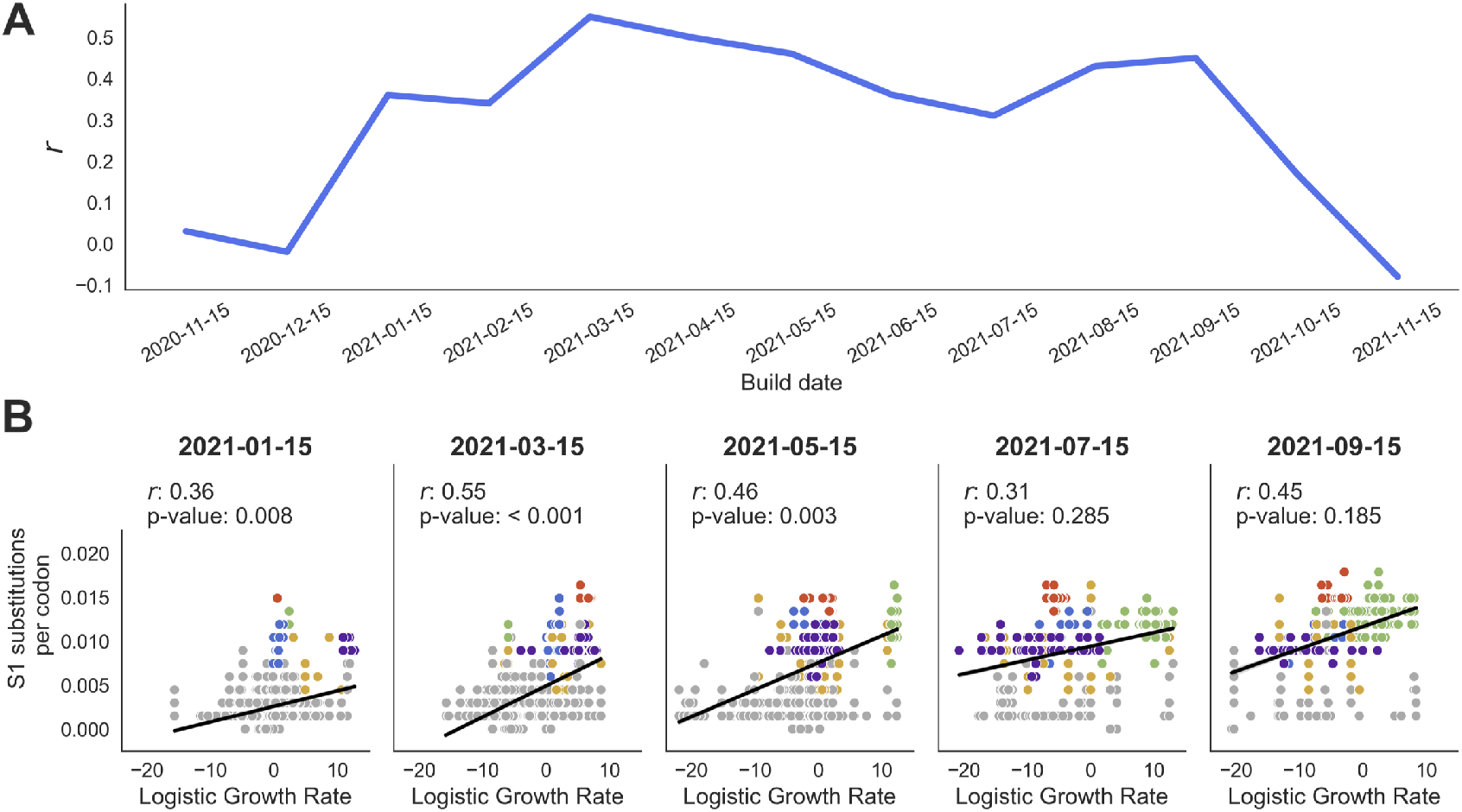
Correlation between S1 substitutions and clade success over time. **A)** The correlation coefficient *r* is calculated between logistic growth rate and S1 substitutions for every clade within 6 weeks preceding the build date (x-axis). **B)** Mutation accumulation is plotted against logistic growth rate and the points are fit by linear regression (as in Figure 1). *p*-values are computed by comparing the observed correlation coefficient *r* to the distribution of *r* coefficients from 1000 trees where S1 substitutions are randomized across branches according to a multinomial draw.

